# A Novel Human TBCK- Neuronal Cell Model Results in Severe Neurodegeneration and Partial Rescue with Mitochondrial Fission Inhibition

**DOI:** 10.1101/2024.10.30.621078

**Authors:** Rajesh Angireddy, Bhanu Chandra Karisetty, Kaitlin A Katsura, Abdias Díaz, Svathi Murali, Sarina Smith, Laura Ohl, Kelly Clark, Andrew V. Kossenkov, Elizabeth J.K. Bhoj

**Affiliations:** Division of Human Genetics, Department of Pediatrics, Children’s Hospital of Philadelphia, Philadelphia, Pennsylvania, PA, USA; Cell and Molecular Biology Graduate Group, Perelman School of Medicine at the University of Pennsylvania, Philadelphia, PA, USA; Department of Orofacial Sciences and Program in Craniofacial Biology, University of California, San Francisco, San Francisco, CA, USA; The Wistar Institute, 3601 Spruce St., Philadelphia, PA 19104

**Keywords:** Neural progenitor cells, mitochondrial dysfunction, cell cycle, neurons, astrocytes, ferroptosis, neurodegeneration

## Abstract

**Background and Objectives:** TBCK syndrome is a rare fatal pediatric neurodegenerative disease caused by biallelic loss-of-function mutations in the *TBCK* gene. Previous studies by our lab and others have implicated mTOR, autophagy, lysosomes, and intracellular mRNA transport, however the exact primary pathologic mechanism is unknown. This gap has prevented the development of targeted therapies.

**Methods:** We employed a human neural progenitor cell line (NPC), ReNcell VM, which can differentiate into neurons and astrocytes, to understand the role of TBCK in mTORC1 activity and neuronal autophagy and cellular mechanisms of pathology. We used shRNA technology to knockdown TBCK in ReNcells.

**Results:** These data showed that loss of TBCK did not inhibit mTORC1 activity in neither NPC nor neurons. Additionally, analysis of eight patient-derived cells and TBCK knock down HeLa cells showed that mTORC1 inhibition is inconsistent across different patients and cell types. We showed that TBCK knockdown in ReNcells affected NPC differentiation to neurons and astrocytes. Specifically, differentiation defects are coupled to cell cycle defects in NPC and increased cell death during differentiation. RNAseq analysis indicated the downregulation of several different neurodevelopmental and differentiation pathways. We observed a higher number of LC3-positive vesicles in the soma and neurites of TBCK knockdown cells. Further, TBCK knockdown altered mitochondrial dynamics and membrane potential in NPC, neurons and astrocytes. We found partial mitochondrial rescue with the mitochondrial fission inhibitor mdivi- 1.

**Discussion:** This work outlines a new Human Cell Model for TBCK-related neurodegeneration and the essential role of mitochondrial health and partial rescue with mitochondrial fission inhibitor. This data, along with human neurons and astrocytes, illuminate mechanisms of neurodegeneration and provide a possible novel therapeutic avenue for affected patients.

## Introduction

TBCK Syndrome, clinically classified as Infantile <u>H</u>ypotonia with <u>P</u>sychomotor <u>R</u>etardation and Characteristic <u>F</u>acies type 3 (IHPRF3), [OMIM 616899], is a rare autosomal recessive disorder that affects brain development in children carrying *TBCK* gene mutations ^1, 2, 3^. This incurable degenerative and aggressive disease is characterized by a wide range of pathologies, including but are not limited to brain atrophy, developmental delay, epilepsy, hypotonia, and respiratory failure ^4^. Although biallelic TBCK mutations are known to have profound implications for brain development, the exact mechanism remains unclear. Understanding this mechanism is essential for developing effective therapies.

<u>TBC</u>1 domain-containing-<u>K</u>inase (TBCK) is a protein encoded by the *TBCK* gene present on chromosome 4 (4q24). The protein contains three different domains: the Protein Kinase domain (PK), Tre-2, Bub2, and Cdc16 domain (TBC), and a Rhodanese homology domain (RHOD). While the functions of kinase domain has been proposed to be catalytically inactive and the rhodanese domain function is unknown, the TBC Rab-GAP domain is predicted to be catalytically active ^5, 6^. TBC domains are evolutionarily conserved from yeast and play an essential role in regulating the activities of small Rab proteins ^7^. Rab proteins are critical members of the vesicle trafficking pathway and coordinate multiple stages of vesicle formation, transport, tethering and organelle motility in all cell types ^8^. Within the last decade, Rab functionality in neurodevelopment and neurodegeneration have become increasingly apparent ^9, 10^. In addition to vesicle transport, TBCK loss modulates mTORC1 activity in patient-derived fibroblasts ^2, 11^.

The <u>m</u>echanistic <u>T</u>arget <u>O</u>f <u>R</u>apamycin (mTOR), a serine/threonine kinase, is a master regulator of brain development in eukaryotic cells. This kinase is responsible for homeostatic functions like protein synthesis, cell proliferation, cell survival, cell migration, and cytoskeleton remodeling ^12^. mTORC can form two multiprotein complexes: mTORC1 and mTORC2. mTORC1 is nutrient- and rapamycin-sensitive and regulates protein synthesis and metabolism. mTORC2 is PI3K- and growth factor-sensitive, but rapamycin-insensitive. The mTORC2 regulates proliferation and autophagy through AKT signaling and also regulates glucose and lipid metabolism ^13, 14, 15^. mTORC1 signaling has been reported to be drastically reduced in cells lacking TBCK fibroblast when they were starved and siRNA knock down in 293T cells ^11^. However, the specific mechanism by which TBCK regulates this pathway is unknown ^2^. Despite this, autophagy is one common pathway affected in TBCK patients ^16^. Accumulation of autophagosomes is associated with poor lysosomal activity. TBCK syndrome is even reported clinically as a lysosomal storage disorder and a neuronal ceroid lipofuscinosis disease ^4, 17^. This clinical classification of NCL reveals a buildup of waste products within cells, resulting in dysfunctional cell death, though the findings varied among autopsy reports. These studies indicate that loss of TBCK affects multiple subcellular organelles and vary by cell type function.

Neurons are highly metabolically active, requiring functional mitochondria to satisfy their high energy demands. Any perturbations in mitochondrial function leads to neurodegenerative diseases ^18, 19^. Recent findings suggest mitochondrial dysfunction is a common phenotype in TBCK syndrome ^17^. In a recent study, TBCK was found to associate with a multi protein complex called FERRY, and this complex play an essential role in mRNA transport for local protein translation. Interestingly, a number of mRNA required for mitochondrial biogenesis was associated with FERRY complex ^20^. Although both mitochondrial and lysosomal defects have been previously reported in TBCK patient-derived fibroblasts, this model cannot mimic the environmental conditions and requirements of the brain. Although one study used NPC cells derived from iPSC and showed that loss of TBCK impaired endoplasmic reticulum-to-Golgi vesicle transport and autophagosome biogenesis and altered cell cycle at NPC, the study lacked isogenic controls ^21^. Thus, to understand the role of TBCK in brain function, we used an immortalized human Neuronal Progenitor Cell (NPC) line ReNcell VM. These NPCs can be readily differentiated into cortical neuronal lineages, including cortical neurons and astrocytes. ^22, 23, 24^. We used shRNA to knockdown TBCK to assess different aspects of the NPCs differentiation and neurobiology. Specifically, we investigated TBCK role in neuronal differentiation, transcription, autophagy, and mitochondrial health. Our results confirm an important regulatory role of TBCK in all of these subcellular processes and offers a new perspective on the protein as a master regulator of neuronal homeostasis.

## Methods

### Cell culture

We used 293T, ReNcell VM, Skin fibroblasts, lymphoblasts, and DF-HeLa cells. All cell lines (except ReNcells) were cultured in DMEM supplemented with 10% fatal bovine serum (FBS) at 37^0^C with 5% CO2. Lymphoblasts were grown as suspension cultures. ReNcells were cultured in N2/B27 medium [DMEM:F12 + Neurobasal medium (1:1 ratio) containing B27 neural cell supplement (Gibco), N2 supplement (Gibco), L-Glutamine (2mM, Gibco), Nonessential amino acids (100µM, Sigma), Insulin (5µg/ml, sigma), betamercaptoethanol (100µM, sigma) and penicillin and streptomycin (100 mM, Gibco)] and expanded on laminin coated (20µg/ml in DMEM/F12 medium) tissue culture plates in the presence of bFGF (20ng/ml, Invitrogen) and EGF (20ng/ml, Sigma), and maintained at 37^0^C in a CO2 incubator. Differentiation was carried out using two different protocols using laminin-coated plates. For standard differentiation, cells were expanded to confluency in growth medium over 2–3 days. Differentiation was initiated by changing to medium lacking growth factors and cultured for two more weeks.

For the pre-aggregation differentiation protocol to generate dopaminergic neurons^22^, cell aggregates were made by seeding 30,000 cells on ultralow attachment 96-well plates (Corning; cat no:12-356-721) in growth medium and expanded for 7 days with media change for alternate days. Aggregates were harvested and dissociated by smaller trituration and replated on laminin-coated 96-well plates. To generate dopaminergic neurons, N2/B27 medium is supplemented with 1 mM dibutyrl-cAMP (Calbiochem) and 2 ng/ml GDNF (Peprotech) to the differentiation media and cultured for seven more days

### Generating stable cell lines

HEK293T cells cultured in 60 mm cell culture dish at 80% confluence were transfected with six different plasmids (pLKO.1-Non-Target shRNA, pLKO.1 TBCK sh1-sh5) obtained from Sigma. shRNA sequences are in Supplementary Table 2. Lentivirus was generated for pLKO.1-Non-Target shRNA, and pLKO.1-TBCK sh5, and concentrated using a kit (Takara; Cat: 631231).

ReNcells were transduced with concentrated virus, and stable cells were generated by treating with 0.4 µg/ml Puromycin. Further, we serially diluted pLKO.1-TBCK sh5 and Scr-Ctrl transduced cells to achieve a one-cell- per-well of a 96-well plate to obtain single cells clones. Similar strategies were applied to generate TBCK KD single-cell clones for DF-HeLa cells.

### RNA sequencing and analysis

Raw reads (FASTQ files) from the RNA sequencing were aligned to the Homo sapiens genome (hg19) using STAR version 2.5.2b ^25^. Gene expression levels were measured using RSEM version 1.3.3 ^26^. Principal component analysis (PCA) was performed in R to cluster the samples and identify the relationships among the samples. Differential gene expression analysis was performed in R using either DESeq2 version 1.38.3 ^27^ or NOISeq version 2.42.0 ^28^; DESeq2 was used for groups with replicates (Differentiated samples), and NOISeq was used for groups with no replicates (Proliferation samples). In NOISeq analysis, simulated replicates for each condition with small variability were generated using default parameters from its manual. For DESeq2, genes with an adjusted P value < 0.05 were considered significant, and for NOISeq, genes with a probability value > 0.9 were considered significant. The probability value, suggested by the NOISeq manual, is not equivalent to the 1 - *P value*. Annotation was done using the EnsDb.Hsapiens.v75 version 2.99.0. The overlapping of significant genes was represented using UpSetR version 1.4.0 ^29^. Gene set enrichment analysis (GSEA) was done for differentiated samples data on R using fgsea version 1.24.0, considering the Enrichr database (https://maayanlab.cloud/Enrichr/#libraries: GO_Biological_Process_2021, GO_Cellular_Component_2021, GO_Molecular_Function_2021, KEGG_2021_Human, Reactome_2022, and MSigDB_Hallmark_2020). The ranked gene list for GSEA analysis was generated by ranking genes using the DESeq2-derived Wald statistic values. Heatmaps with the genes of selected pathways were generated using Complex Heatmap version 2.14.0 ^30^. All other plots were constructed using ggplot2 version 3.4.0.

### Neurosphere assay

To make equal sized neurospheres, we used AggreWell-800 plates (Stemcell Technologies; Cat: 34815). AggreWell plates were rinsed with 500µl anti-adherence solution and centrifuged at 1300xg for 5min to remove air bubbles. The rinsing solution was replaced with 1ml of B2/N27 medium with growth factors. Then added 3×10^6^ cells/ mL into each well (10,000 cells/microwell), and centrifuged the plate at 100 × g for 3 min to capture cells in the microwells. Cells were incubated at 37^0^C in a CO2 incubator for six days with partial medium changes on alternate days (Stem Cell Technologies technical manual). Neurosphere images were taken using an Evos XL Core light microscope at 10X magnification and measured using Image J.

### Immunocytochemistry

Cells were fixed in 4% paraformaldehyde, washed three times with ice-cold PBS, permeabilized with 0.1% Triton X-100 for 15 min, and blocked with 5% goat serum in PBS for one hour at 25°C. Cells were incubated with primary antibody overnight at 4^0^C. After washing with PBSx3, the cells were incubated with Alexa fluor-conjugated secondary antibodies for another hour at 25°C. All antibodies and dilutions are reported in supplementary table 3. After three PBS washes, cells were stained with DAPI and mounted on glass slides using Fluoromount-G^TM^ (ThermoFischer Scientific). For MitoTracker™ Orange CMTMRos staining, cells were treated with 200nM dye for 45min and washed three time with PBS and fixed in 4% PFA. Immunofluorescence was done as mentioned above. Images were taken on Leica-SP8 confocal or Keyence fluorescence microscopes at different magnifications. Fluorescence intensity was calculated using Image-J and expressed as corrected total cell fluorescence (CTCF), calculated by subtracting the integrated density value from the area of the selected cell multiplied by the mean fluorescence of background.

### Western blot analysis

Cells were homogenized in RIPA buffer (final concentrations: 50 mM Tris, 150 mM NaCl, 1% Triton-X100, 0.1% SDS, 0.5% sodium deoxycholate, with Complete protease and phosphatase inhibitors (Roche), and samples were then clarified by centrifugation at 10,000g for 10min. Total protein concentration was determined by BCA assay (Thermo Scientific; Cat: 23225), and 10μg of total protein was loaded onto 4–12% NuPAGE Bis-tris gels in MES buffer (Invitrogen; Cat:NP0323). After electrophoresis, proteins were transferred to 0.45 μm PVDF (Thermo Scientific; Cat: 88518). Membranes were blocked with 2% BSA-PBS (for detecting phosphorylated proteins) or 5% milk and incubated with different primary antibodies dissolved in 2% BSA-PBS, followed by horseradish-peroxidase-coupled secondary antibodies. Blots were developed with ECL Plus reagents (Thermo Scientific; Cat: 34577) and imaged on the ChemiDoc Imaging System (BioRad). Band intensities were calculated using image J. All antibodies and dilutions are presented in Supplemental Table 3.

### Gene expression analysis

Cells were dissolved in TRIzol (Life Technologies; Cat: 15596018), and RNA was isolated using Direct-zol RNA Miniprep Kit (Zymo Research; Cat: R2050) according to the manufacturer’s instructions. 1µg of total RNA was reverse-transcribed using SuperScript™ VILO MasterMix (ThermoFischer Scientific; Cat:11755050) according to the manufacturer’s protocol. RT-qPCR analysis was performed in Quant studio-3 PCR System using a Power-SYBR Green master mix (Applied Biosystems; Cat:100029284). Primers used in the study are shown in Supplementary Table 2. Relative mRNA levels were calculated using 2^−ΔΔCt^ method using GAPDH as a reference gene.

### BrdU assay

Scr-Ctrl and TBCK KD ReNcells cells were seeded on a 24-well plate (3×10^3^ cells/well) with 12mm PDL coverslips coated with laminin in N2/B27 medium. The next day, 3 µg/ml BrdU (Invitrogen; Cat:00-0103) was added and incubated overnight. For BrdU staining, the cells were fixed with 4% paraformaldehyde, washed thrice with PBS, and treated with 2N HCL for 30min and washed three times with PBS. Cells were incubated in 0.1% Triton X-100 in PBS for 20min at 25°C and washed thrice with PBS. Then cells were blocked with 5% goat serum, treated with Anti-BrdU antibody, and continued with the same immunofluoresecence protocol described above. Following imaging, BrdU positive cells were manually counted, and the percentage of BrdU positive cells from the total population was calculated.

### Cell cycle analysis by Propidium Iodide staining

For this assay, we used FxCycle™ PI/RNase Staining Solution (F10797, Invitrogen), which comes with DNase-free RNase A and a permeabilization reagent to accurately stain the DNA content of cells. Measuring the DNA content by flow cytometry is a standard method to understand various phases of cell cycles in a given population. For this study, proliferating ReNcells were plated on a 6-well culture plate cultured for 24hr. Flow cytometry was carried out to analyze the cellular DNA content. Fluorescence intensity of stained nuclei was measured with a flow cytometer (BD FACS Calibur), and data were analyzed using FlowJo software.

### ROS analysis

We measured Hydrogen peroxide, superoxide in differentiated cells using two methods. To measure H2O2, we used AmplexRed Assay, where H2O2 reacts with Aplexred in the presence of horseradish peroxidase (HRP) and forms the red fluorescent oxidation product Resourfin. 30,000 ReNcells were plated on a 96-well plate and differentiated into neurons and astrocytes. Cells were treated with AmplexRed (50μM), HRP (0.2U) and incubated for 30min at 37°C. The resulting fluorescence was measured using a fluorescent microplate reader at 530nm^ex^/590nm^em^ (BioTek, USA). Fluorescent readings were normalized to total protein. Similarly, wemeasured superoxide using ROS-ID^®^ Total ROS/Superoxide detection kit (Enzo Lifesciences). After plating, however, cells were treated with 2µM superoxide detection reagent and incubated for 60min at 37°C in CO2 incubator. Fluorescenc was measured at 550nm^ex^/620nm^em^ (BioTek, USA) and normalized.

### Autophagy analysis in DF-HeLa cells

Scr-Ctrl and TBCK KD DH-HeLA^31^ cells were cultured in DMEM with 10% FBS containing Normocin (100µg/ml) and Zeocin (100µg/ml). For microscopy, 30,000 cells/well were plated on chamber slides, cultured for 24hr, fixed with 4% PFA, then permeabilized with 0.1% Triton X100 and stained with DAPI. For protein expression using western blot, 1×10^6^ cells were plated on 60mm plates and total cell lysates were prepared using RIPA buffer. For induction of autophagy, cells were treated overnight with 100nM rapamycin.

### Statistical analysis

All experiments were performed in biological triplicates. Multiple comparisons were done using one-way ANOVA followed by a Bonferroni’s post-hoc test (Graph Pad PRISM). Student’s t-test was used to calculate statistical significance for comparing two groups (Graph Pad PRISM). A p-value of less than 0.05 was considered significant (*p-value < 0.05, **p-value < 0.01, ***p-value < 0.001). Error bars represent the mean ± standard error of the mean.

### Standard Protocol Approvals, Registrations, and Patient Consents

This study does not include any human subjects research or animal models, there are no recognizable persons included and not clinical trial data.

## Results

### mTORC1 activity varies in TBCK-knockdown ReNcells and patient-derived cells

Children with TBCK syndrome show severe neurodegeneration leading to seizures, neurologic decline, central respiratory insufficiency, and early death ^2, 4, 16^. To understand the effects of loss of TBCK in neuronal differentiation and function *in vitro*, we used a human neural progenitor cell line (ReNcell VM), which readily differentiated into neurons and astrocytes ^22^. As previously reported, ReNcells differentiate into neurons and astrocytes after culturing in N2/B27 medium without growth factors for two weeks **(Supplemental Figure. 1A)**. To knockdown TBCK using shRNA, we transiently transfected 293T cells with either the pLKO.1 non-targeting scramble control (Scr-Ctrl) or one of five different shRNA that targeted different exons of the TBCK mRNA (pLKO.1 TBCK sh1-sh5). We observed that sh4 and sh5 had the maximum knockdown efficiency in 293T cells **(Supplementary** Figure. 1B). After confirming knockdown efficiency in 293T cells, we transduced ReNcells with different dilutions of lentivirus carrying the pLKO.1 Scr-Ctrl or pLKO.1 TBCK SH-5. Cells transduced with the TBCK SH-5 reduced TBCK mRNA levels in a dose-independent manner **(Supplemental Figure. 1C)**. Single-cell clones were isolated from transduced cells, and qRT-PCR analysis indicated that most single-cell clones had knockdown efficiency between 70-80% **(Figure. 1A & Supplementary** Figure. 1D). Single cell clones (TBCK KD-sc1, TBCK KD-sc2) and single cell clone from Scr-Ctrl were used for further analysis.

**Figure 1.**
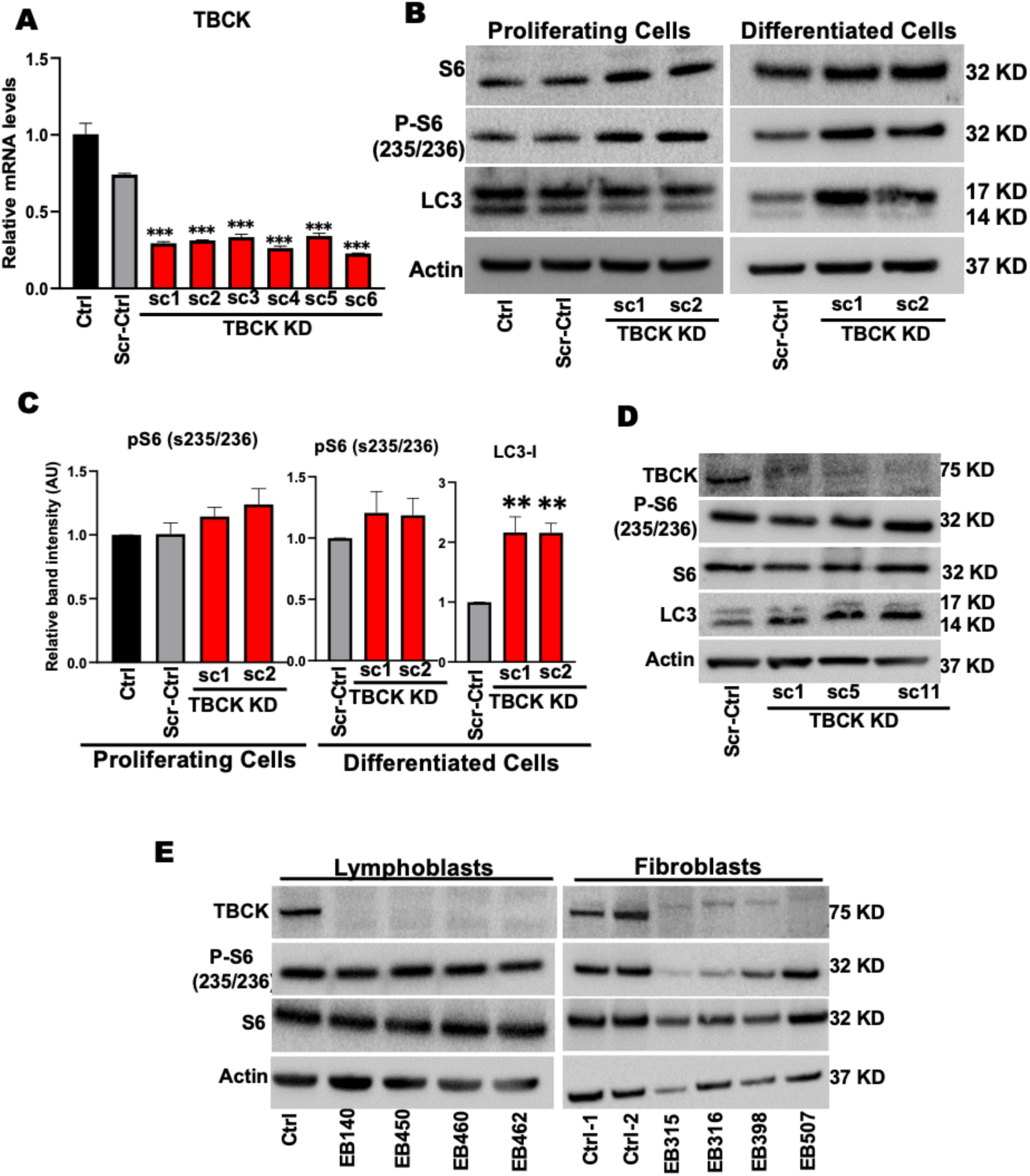
TBCK knockdown in ReNcells do not down regulate mTORC1. **A:** qRT-PCR analysis of TBCK mRNA levels in lentivirus transduced single cell clones. **B:** Representative Western blots of proliferating and differentiated ReNcells for Control, Scr-Ctrl, TBCK KD sc-1, and TBCK KD sc-2 for S6, p-S6, LC3, and Actin done in triplicate. **C:** Band intensities were calculated in Image J normalized to Actin. The relative band intensity compared to control was presented. Error bars represent SEM. *vs. Scr-Ctrl. **D:** Representative Western blots of Control and TBCK knockdown DF-HeLa cells for TBCK, S6, p-S6, LC3, and Actin done in triplicate. **E:** Representative Western blots of controls and patient-derived lymphoblasts and fibroblast for S6, p-S6 and Actin done in triplicate.

Studies from patient-derived fibroblasts indicate that loss of TBCK inhibits mTORC1 activity ^2^ and increases accumulation of autophagosomes, illustrated by LC3 immunostaining ^16^. To investigate whether this is the case in NPC and neurons, we measured p-S6 and LC3 protein levels in TBCK KD, ReNcells. Interestingly, knockdown of TBCK in proliferating and differentiated ReNcells had no effect on phosphorylation of S6. However, we observed slightly higher phosphorylation of S6 in two different TBCK knockdown sub clones **(Figure. 1B & C)**. LC3 levels were comparable to control in proliferating TBCK KD ReNcells (**Figure. 1B**), but we observed higher LC3 in differentiated TBCK KD cells **(Figure. 1B & C)**. When we directly compared the mTORC1 activity between proliferating and differentiated ReNcells **(Supplemental Figure. 1E)**, we observed significantly lower mTORC1 activity in differentiated ReNcells regardless of the genotype (pS6 as a proxy) in differentiated cells. Low mTORC1 activity in differentiated cells may be a compounding factor for higher LC3 levels in TBCK KD differentiated cells.

To tease out if these trends corresponded to other cell types, we used different clones of TBCK knockdown DF-HeLa cells to look at p-S6 and LC3 levels **(Supplemental Figure. 1F)**. These HeLa cells express LC3 protein conjugated with GFP and RFP to differentiate autophagosome vs autolysosome accumulation ^31^. We observed no difference in p-S6 levels, but there was a significant increase in LC3-II in different TBCK KD clones **(Figure. 1D & Supplemental Figure. 1G)**. Since TBCK knockdown in ReNcells and DF-HeLa did not inhibit mTORC1 signaling, we analyzed p-S6 levels in patient-derived TBCK fibroblasts and lymphoblasts **(Supplemental Table. 1)**. Patient-derived TBCK lymphoblasts carrying different pathogenic variants, there were no differences in p-S6 levels. Whereas dermal fibroblasts with the homozygous loss of function mutation had low levels of both total S6 and p-S6 levels, indicating total pathway reduction **(Figure. 1E & Supplemental Figure. 2A)**. These results indicate that in the absence of TBCK, mTORC1 signaling is not always downregulated. Instead, our results suggest that mTORC1 activation is variable with reduced TBCK levels, and is a process heavily modulated by cell type and mutation.

### TBCK knockdown affects ReNcell differentiation into neurons and astrocytes

To understand the effects of TBCK knockdown on differentiation, Scr-Ctrl and TBCK KD NPCs were differentiated into neurons and astrocytes **(Figure 2A)**. After two weeks of differentiation, TBCK KD neuronal somas had reduced cluster formation, and reduced neuronal projections, as compared to the Scr-Ctrl cells, which showed cell aggregation indicated by clustered somas with neuronal and glial projections in organized co-tract-like formations, as observed by TUJ1 and GFAP staining **(Figure 2B, Supplemental Figure. 2B)**. Similar results were also observed in an additional TBCK KD monoclonal subpopulation **(Supplemental Figure. 2C)**. Intensity of TUJ1, MAP2 and GFAP markers were also decreased in TBCK KD differentiated cells **(Figure 2B, Supplemental Figure. 2D & E)**, which accompanied significantly reduced MAP2 and GFAP mRNA levels **(Figure. 2C)**. Expression of a neural progenitor marker (PAX6) appeared only in NPC and slightly elevated in TBCK KD cells at the proliferating stage, while the neuronal maturation marker (MAP2) was significantly lower in TBCK KD cells at the differentiated stage compared to Scr-Ctl **(Figure. 2D & Supplemental Figure. 2F)**. Conversely, we saw increased GFAP in differentiated TBCK KD cells (**Figure 2B**, **2D & Supplemental Figure 2F**). We stained differentiated ReNcells with another neuronal maturation marker, NeuN, and found a significant reduction in the number of NeuN positive soma in TBCK KD cells **(Figure. 2E)**. These results revealed that loss of TBCK delays or prevents neuronal differentiation and maturation, in addition to neurite formation.

**Figure 2.**
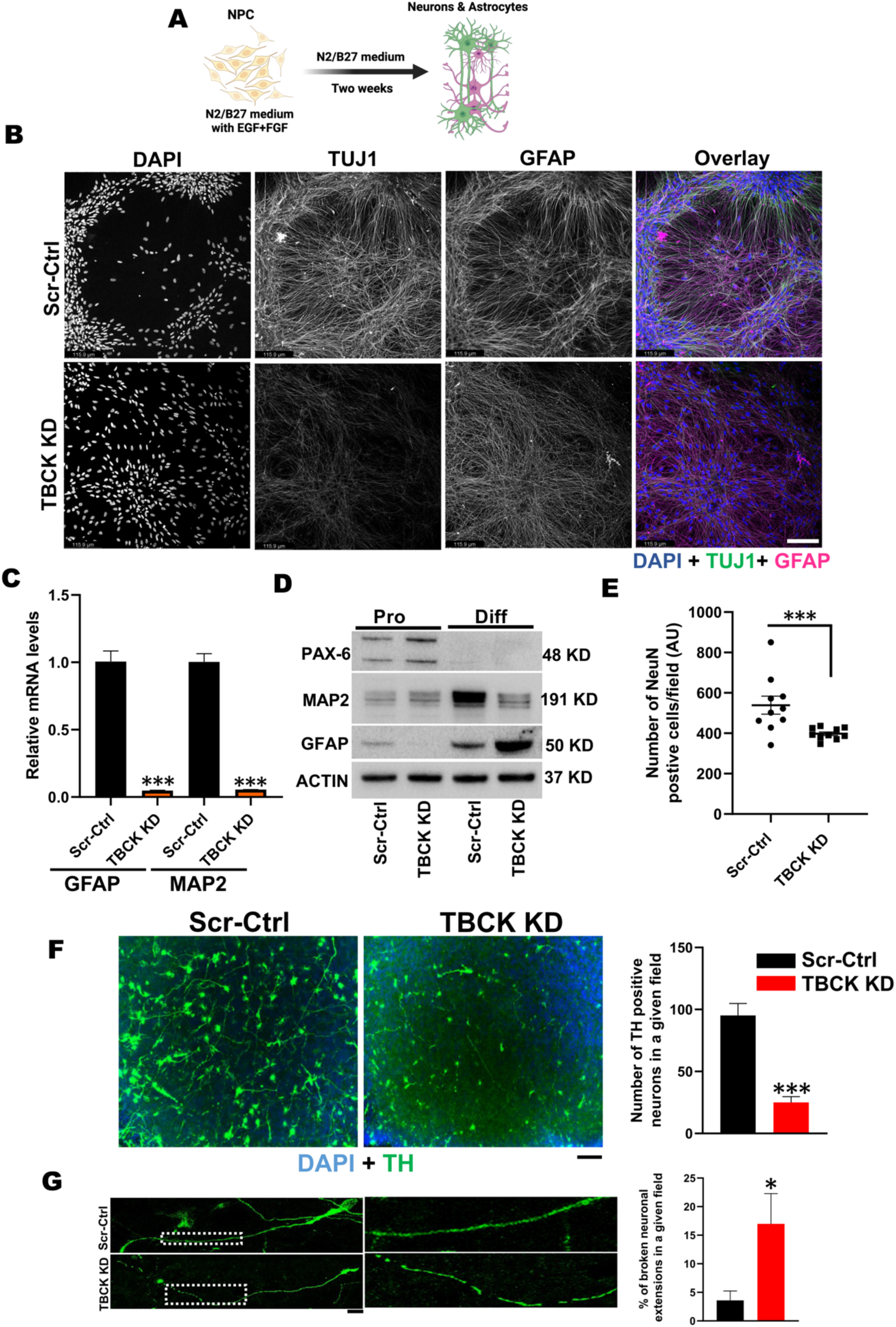
TBCK knockdown in ReNcells affected their neuronal and glial differentiation. **A:** Pictorial representation of ReNcell differentiation into neurons and astrocytes. **B:** TUJ1, GFAP, and DAPI immunofluorescence staining for differentiated Scr-Ctrl and TBCK KD ReNcells done in triplicate. Images were taken at 20X magnification, scale bar represents 115.9µm. **C:** qRT-PCR analysis of MAP2 and GFAP mRNA levels in differentiated Scr-Ctrl and TBCK KD ReNcells. Error bars represent SEM. *vs. Scr-Ctrl **D:** Representative Western blots of proliferating and differentiated ReNcells for PAX6, MAP2, GFAP, and Actin done in triplicate. **E:** Immunofluorescence staining of NeuN followed by counting the number of NeuN-positive cells in a given field. Data is the average of three experiments. **F:** Immunofluorescence staining of Scr-Ctrl and TBCK KD ReNcells differentiated to dopaminergic neurons for Tyrosine Hydroxylase (TH) and DAPI. Images were taken at 20X magnification. The number of TH positive cells in a given field is presented as a graph. Scale bar is 10µm. Error bars represent SEM. *vs. Scr-Ctrl, **G:** Enlarged image showing TH positive neuronal extensions. The percentage of broken extensions in a given field is presented as a graph. Scale bar is 50µm. Error bars represent SEM. *vs. Scr-Ctrl

In addition to dysfunction of cortical neurons being implicated in intellectual disability, TBCK patients also have motor impairment and many patients never achieve independent walking, brain atrophy, corpus callosum dysgenesis, and cerebellar vermis hypoplasia ^2, 4^. Dopaminergic (DA) neurons of the midbrain regulate voluntary movement, ^32^ and loss of these neurons is associated with Parkinson’s disease (PD) and Alzheimer’s disease ^33, 34^. Since the degenerative clinical symptoms in TBCK patients mimic other neurodegenerative diseases, we wanted to investigate if TBCK also plays a role in the differentiation into TH-positive dopaminergic neurons. Using TH as a marker, we observed not only a significant defect in the differentiation of DA neurons but also a significant number of breaks along the length of neuronal extensions in these cells **(Figure 2F & G)**. Together with our cortical and DA neuron differentiation findings reveal that knockdown of TBCK affects ReNcell differentiation into different neuronal subtypes.

### TBCK knockdown induced proliferation defects in NPCs and pro-apoptotic protein induction during differentiation

Due to the severe defects in TBCK KD ReNcells in NPC differentiation to neurons and astrocytes, we wanted to understand the possible mechanism for this reduced differentiation. To do this we investigated if cell proliferation was impacted and if cell death was induced. To determine whether cell proliferation was affected, we stained proliferating cells with BrdU and observed significantly fewer BrdU positive cells in TBCK KD cells **(Figure. 3A)**. Further analyses using a 3D neurosphere assay also showed proliferation defects in TBCK KD cells. While we observed significant increases in neurosphere size in Scr-Ctrl cells, TBCK KD cells failed to grow **(Figure. 3B & C)**. Further assessment of the cell cycle profiles showed a higher number of cells in the G0/G1 stage and fewer in the G2/M cell stage in the TBCK KD proliferating cells, indicative of increased quiescent or senescence cells **(Figure. 3D)**. Additionally, TBCK KD differentiated cells showed reduced levels of anti-apoptotic marker, BCL-2, and higher levels of pro-apoptotic markers BID, BAX and caspase-9 **(Figure 3E & Supplemental Figure. 3A & B)**. While caspase- 3 levels were increased in differentiated cells irrespective of genotypes compared to proliferating cells, we could not detect cleaved caspase-3 levels **(Figure. 3E & Supplemental Figure. 3B)**. These results indicate that TBCK knockdown affects cell cycles in the proliferating NPC cells and induces apoptosis during differentiation.

**Figure 3.**
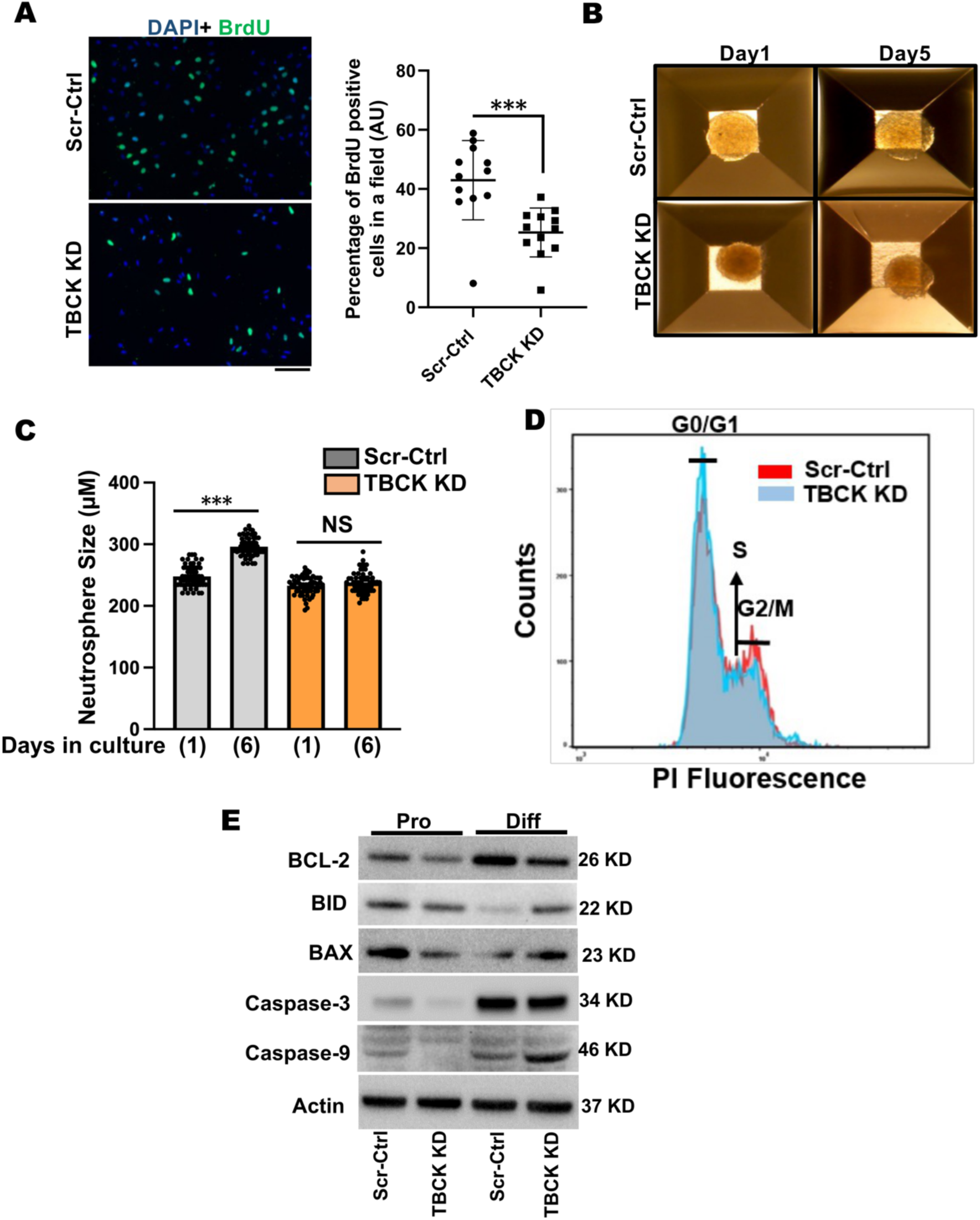
TBCK knockdown affected cell proliferation in ReNcells. **A:** BrdU staining of proliferating Scr-Ctrl and TBCK KD ReNcells. Images were taken at 40X magnification. The number of BrdU-positive cells in a given field is counted and plotted as a percentage change. Data is the average of three experiments. Error bars represent SEM. *vs. Scr-Ctrl. **B & C:** Equal-sized neurospheres are made in Aggrewell 800 plates, and size was calculated using Image J. Data is the average of three experiments. Error bars represent SEM. *vs. Scr-Ctrl. **D:** Scr-Ctrl and TBCK KD proliferating ReNcells were stained with propidium iodide and cell cycle progression is analyzed by flow cytometry. Pictorial graph showing the proportion of cells in different phases of the cell cycle. **E:** Representative Western blots of proliferating and differentiated ReNcells for BCL-2, BID, BAX, Caspase3, Caspase9, and Actin done in triplicate.

### TBCK knockdown affects the transcriptome of proliferating and differentiated ReNcells

We performed bulk RNA sequencing on both proliferating and differentiated TBCK KD cells to understand the pathways affected at these two stages of neurodevelopment. PCA analysis showed the difference between proliferating and differentiated cells on the principal component 1 (PC1) and a considerable difference between the Scr-Ctrl and TBCK KD groups on the PC2 **(Supplementary** Figure. 3C). We used NOISeq and DESeq2 pipelines to identify TBCK KD-induced changes in gene expression profile (TBCK KD vs. Scr-Ctrl). Although TBCK KD had lesser effect on the gene expression (73 genes significantly upregulated and 112 genes significantly downregulated) in proliferating cells **(Figure. 4A & Supplementary** Figure. 3D), we observed a sizeable number of differentially regulated transcripts in differentiated cells (1303 transcripts significantly upregulated and 1072 transcripts significantly downregulated) **(Figure. 4B & Supplementary** Figure. 3D). Overlapping of the TBCK KD-induced differentially regulated gene transcript from proliferating and differentiated cells revealed that 42% of transcripts were upregulated (47 out of 112) and 52% of downregulated (38 out of 73) **(Supplementary** Figure. 3D). We have presented all the genes and pathways significantly up or down regulated in the proliferating and differentiated cells in **supplemental worksheet**. Our bulk RNAseq results implicate inherent defects in NPCs that prevent proper differentiation into neural and glial lineages. Additionally, this signifies that additional pathways are impacted by TBCK knockdown after differentiation into neurons and astrocyte populations.

**Figure 4.**
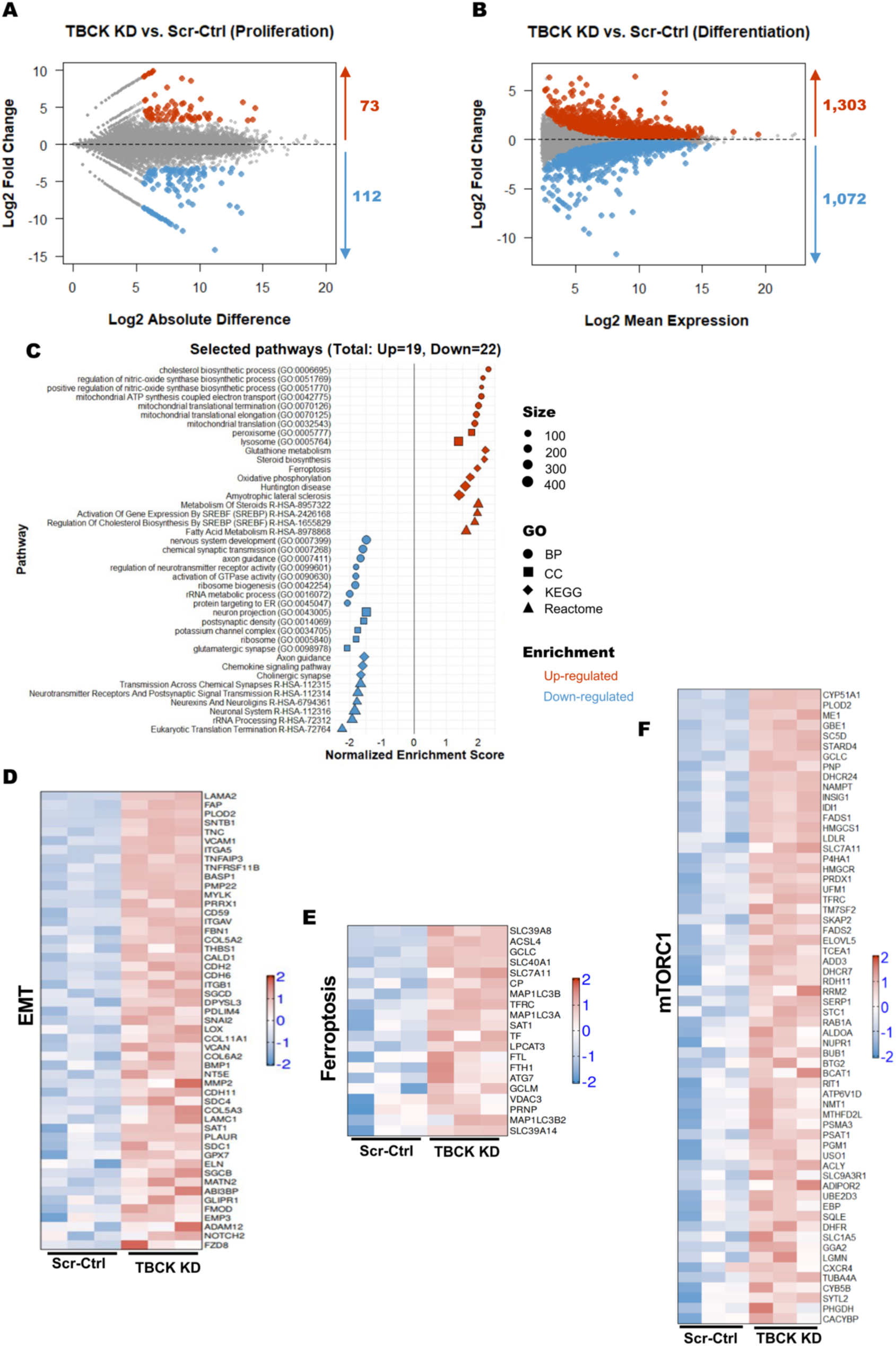
RNASeq analysis of Scr-Ctrl and TBCK KD ReNcells reveled different pathways for neurodegeneration. **A:** MD plot of RNA-seq expression of 25,990 genes between Scr-Ctrl proliferating and TBCK KD proliferating cells. Red dots indicate differentially expressed genes that are upregulated in TBCK KD proliferating cells (Probability value > 0.9 and log2 fold change > 0), while blue dots indicate differentially expressed genes that are downregulated in TBCK KD proliferating cells (Probability value > 0.9 and log2 fold change < 0). **B:** MA plot of RNA-seq expression of 57,736 genes between Scr-Ctrl differentiated and TBCK KD differentiated cells. Red dots indicate differentially expressed genes that are upregulated in TBCK KD differentiated cells (adjusted P value < 0.05 and log2 fold change > 0), while blue dots indicate differentially expressed genes that are downregulated in TBCK KD differentiated cells (adjusted P value < 0.05 and log2 fold change < 0). **C:** GSEA was performed using different gene sets from Enrichr database with the significantly regulated genes from differentiated cells (TBCK KD vs. Scr-Ctrl). Selected pathways from different gene sets (Biological Process, BP; Cellular Component, CC; KEGG; Reactome) with adjusted P values (padj <0.05) are shown. The shape represents the source of different gene sets; color represents the direction of change; size represents the gene set size. **D to F:** Heatmap representation of gene expression from selected pathways: **(D)** Epithelial Mesenchymal Transition, MSigDB hallmark gene set; **(E)** Ferroptosis, KEGG; and **(F)** mTORC1 signaling, MSigDB hallmark gene set. Log-transformed gene expression values are displayed as colors ranging from red to blue, as shown in the key. Red represents an increase in gene expression, while blue represents a decrease in expression.

Therefore, we focused on differentiated ReNcells for gene ontology and pathway analysis using GSEA. We observed several pathways to be significantly altered. Critical pathways that significantly upregulated were ferroptosis, cholesterol and sterol biosynthesis, epithelial-mesenchymal transition (EMT), and different mitochondrial pathways **(Figure. 4C)**. There were also a group of pathways associated with neuronal health, signaling and synopsis, and calcium and potassium channels that were significantly downregulated **(Figure. 4C)**. Beyond neuronal-specific processes, ribosome and protein translation pathways were also downregulated. Increased expression of several genes associated with EMT and ferroptosis pathways implicate that these two pathways may play an essential role in the differentiation and cell survival defects we observed **(Figure 4D & E)**. Likewise, many mTORC1 signaling associated genes were significantly upregulated in differentiated cells **(Figure. 4F)**. Overall, these results show that many shared genes are affected in TBCK KD proliferating and differentiated cells. Unsurprisingly, based on our previous results, differentiated cells lacking TBCK have upregulation of several pathways associated with neurodegeneration.

Since our bulk RNA-Seq analysis revealed ferroptosis induction during differentiation of TBCK KD cells, we wanted to investigate if this was mediated by reactive oxygen species (ROS), which play an essential role in the induction of ferroptosis ^35^. We measured H2O2 and O2^-^ and observed that both were significantly increased in differentiated TBCK KD ReNcells **(Figure. 5A & B)**. To further validate upregulation of ferroptosis in differentiated TBCK KD ReNcells, we measured a marker of ferroptosis, GPX4 ^35^. Differentiated TBCK KD cells had significantly lower levels of the GPX4 protein **(Figure. 5C & Supplemental Figure. 3E)**, indicating that TBCK KD increases ROS, initiates ferroptosis, and induces apoptosis programming leading to cell death in TBCK KD differentiated cells.

**Figure 5.**
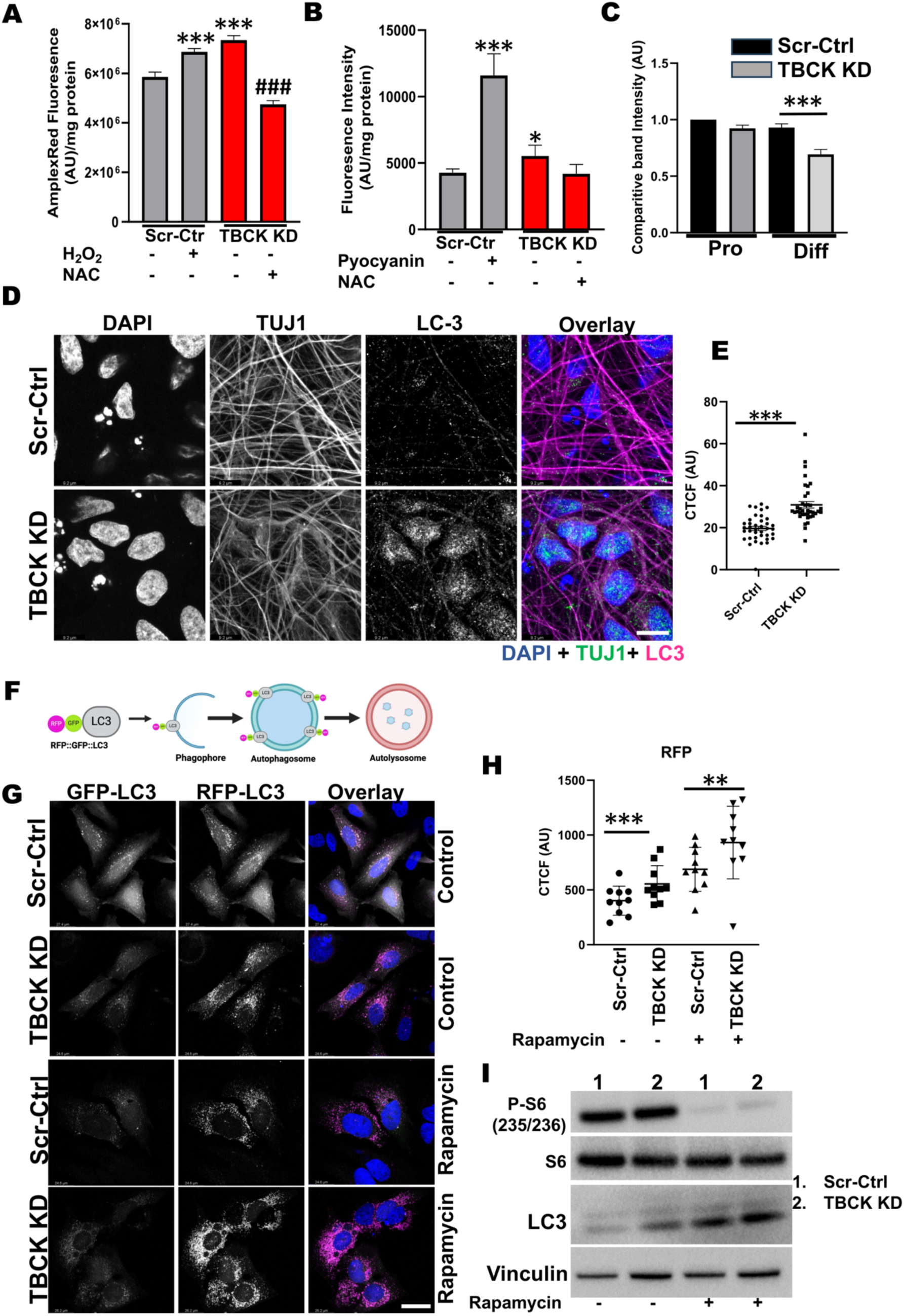
Differentiated ReNcells with TBCK knockdown has increased ROS and autophagy. A: ROS measurements in RenCells differentiated into neurons and astrocytes by AmplexRed assay. Scr-Ctrl cells were treated with 20 µM hydrogen peroxide as a positive control. TBCK KD cells were treated with 5mM NAC as an assay specific control. Error bars represent SEM. ***vs. Scr-Ctrl. ### vs. TBCK KD, **B:** Superoxide levels in RenCells differentiated into neurons and astrocytes were measured using ROS- ID^®^ Superoxide detection reagent. Scr-Ctrl cells were treated with 50 µM pyocyanin as a positive control. TBCK KD cells were treated with 5mM NAC as an assay specific control. Error bars represent SEM. *vs. Scr-Ctrl. **C:** Western blot quantifications of proliferating and differentiated ReNcells for GPX4 and Actin. Band intensities were calculated using Image J normalized to Actin. The relative band intensity compared to control was presented. Error bars represent SEM. ***vs. Scr-Ctrl **D:** Immunofluorescence staining of LC3, TUJ1, and DAPI for differentiated Scr-Ctrl and TBCK KD ReNcells. Images were taken at 63X magnification, scale bar = 9.2µm. **E:** The fluorescence intensity of the soma region is plotted as a graph. Error bars represent SEM. ***vs. Scr-Ctrl. **F:** Pictorial representation of DF-HeLa cells expressing LC3 conjugated with GFP and RFP. When autophagosomes fuses with lysosomes, GFP is quenched, expressing only RPP conjugated LC3. **G:** Scr-Ctrl and TBCK KD DF-HeLA cells treated with rapamycin and accumulation of GFP and RFP puncta were imaged at 63X magnification, scale bar = 24.6µm. **H:** The RFP intensities of the cells were calculated using image J and expressed as corrected total cell fluorescence (CTCF). Error bars represent SEM. *** vs. Scr-Ctrl. **I:** Represesentative Western blots of Scr-Ctrl and TBCK KD DF-HeLa treated with 100nM Rapamycin for S6, pS6, LC3 and Actin done in triplicate. All the immunofluorescence images represent three independent experiments.

### Differentiated TBCK KD cells show increased autophagy and accumulation of transport markers in dead cells

Although we found evidence to suggest apoptosis was being induced due to reduce TBCK levels, the question as to how this was being mediated upstream remained. Therefore, we next investigated the role of TBCK in neuronal autophagy. First, we stained Scr-Ctrl and TBCK KD differentiated cells with a neuronal marker (TUJ1) and autophagy marker (LC3). We observed accumulation of LC3-positive vesicles in the soma of TBCK KD cells compared to Scr-Ctrl **(Figure. 5D & E)** and increased LC3-positive vesicles in neuronal projections of TBCK KD cells **(Supplementary** Figure. 3F). In order to distinguish whether these LC3-positive vesicles were autophagosomes or autolysosomes, we used a reporter line containing LC3-conjugated with GFP and RFP (DF-HeLa). When autophagosomes fuse with the lysosomes, GFP is quenched inside the lysosomes, while RFP remains stable **(Figure. 5F)**. In TBCK KD DF-HeLa cells at baseline we observed higher RFP vesicles compared to Scr-Ctrl **(Figure. 5G & H)** but GFP vesicles remained the same **(Figure. 5H & Supplementary** Figure. 3G). When treated with rapamycin, a repressor of mTOR signaling, we observed significantly higher RFP in TBCK KD cells compared to Scr-Ctrl cells **(Figure. 5G, H &I)**. We observed similar increases in LC3-II levels from western blot analysis indicating accumulation of matured autophagosome accumulation. **(Figure. 5I & Supplemental Figure. 4A)**. These results implicate that autophagy induction is not affected in TBCK KD DF-HeLa cells but the degradation of autophagosomes inside the lysosomes are affected. In summary, loss of TBCK results in accumulation of autophagosomes in both ReNcell neurons and DF-HeLa cells. Additionally, with the DF-HeLa we prove that the higher LC3 level in TBCK KD cells is the result of incomplete degradation of autophagosomes inside the lysosomes.

Our bulk RNAseq analyses on differentiated cells indicated significant down-regulation of genes that play an important role in synaptic transmission and axonal guidance **(Figure. 4C)**. Synpasin-1 plays an important role in nerve conduction at the axon terminals, so to understand the distribution of Synapsin-1, we stained differentiated ReNcells with SYN1 and co-stained with neurite marker (TUJ1). Surprisingly, we observed significantly higher numbers of Syn-1-positive cells with fragmented nuclei in the TBCK KD differentiated cells, the latter indicating that they were dead **(Figure. 6A & B)**. Synapsin1 is produced in the soma and transported to axon terminals in mature neurons ^36^. Presence of Syn-1-positive cells with fragmented nuclei implicates that either SYN1 accumulates in the soma due to a lack of projections forming, or due to impaired axonal transport leading to cell death.

**Figure 6.**
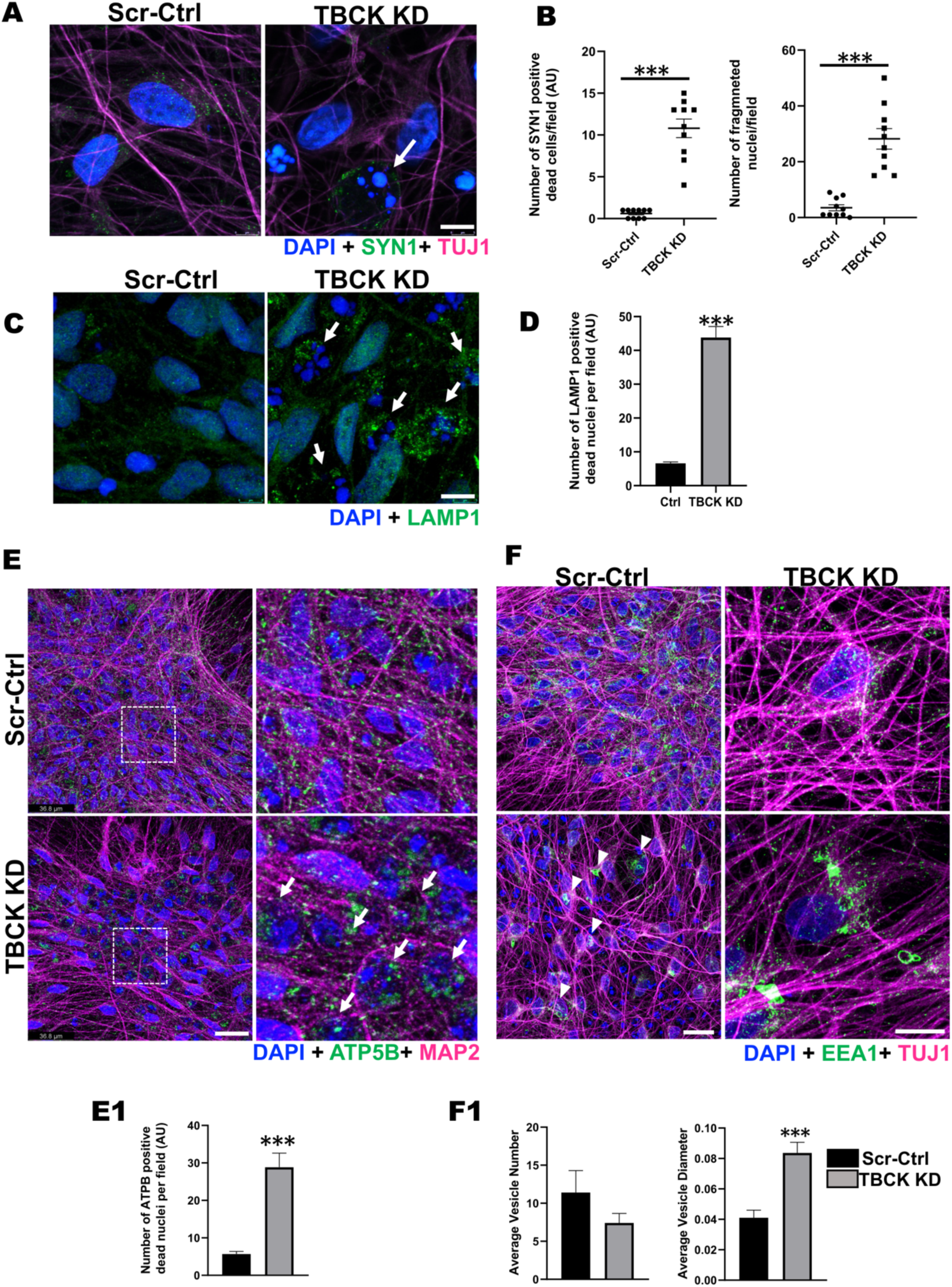
TBCK KD differentiated ReNcells accumulated dead cells staining for different transport proteins. **A & B: (A)** Immunofluorescence staining of SYN1, TUJ1, and DAPI for differentiated Scr-Ctrl and TBCK KD ReNcells. Images were taken at 63X magnification, scale bar = 7µm. **(B)** The number of dead cells and dead cells positive for SYN1 is presented as a graph. Error bars represent SEM. *** vs. Scr-Ctrl. **C & D: (C)** Immunofluorescence staining of LAMP1 and DAPI for differentiated Scr-Ctrl and TBCK KD ReNcells. Images were taken at 63X magnification, scale bar = 7µm. **(D)** The number of dead cells positive for LAMP1 is presented as a graph. Data represent an average of three experiments. Error bars represent SEM. *** vs. Scr-Ctrl. **E:** Immunofluorescence staining of ATP5B, MAP2, and DAPI for differentiated Scr-Ctrl and TBCK KD ReNcells. Images were taken at 63X magnification, scale bar = 36.8µm. Panels on the right show equally zoomed in views of panels on the left outlined in dashed-white boxes. White arrows denote accumulated ATP5B. **(E1)** The number of dead cells positive for ATP5B is presented as a graph. Error bars represent SEM. *** vs. Scr-Ctrl **F:** Immunofluorescence staining of EEA1, TUJ1, and DAPI for differentiated Scr-Ctrl and TBCK KD ReNcells. Images were taken at 63X magnification, scale bar = 50µm. **(F1)** Average number of vesicles in the soma and the average diameter of the vesicles are presented as a graph. Error bars represent SEM. *** vs. Scr-Ctrl. All the immunofluorescence images represent three independent experiments.

To further understand if dead cells are positive for other organelles which are actively transported across the neurons was also impaired due to TBCK knockdown, we stained for lysosomes (LAMP1) and mitochondria (ATP5B). In TBCK KD differentiated cells, we observed LAMP1+ and ATP5B+ cells with fragmented nuclei in significantly higher numbers **(Figure. 6C, D, E & E1)**. Therefore, we also investigated if the transport of vesicles was impacted, such as early endosomal pathways that transport multiple cargoes including mRNAs for local protein translation ^20^. We stained ReNcells for early endosomes (EEA1) and observed a decreased number of endocytic vesicles per soma in TBCK KD cells, but a significant increase in their diameters **(Figure. 6F & F1)**. The presence of significantly larger vesicles implies fusion of multiple vesicles due to accumulation in soma or improper transport. These results indicate that lack of TBCK affects the distribution of different transport vesicles (early endosomes), organelles (lysosomes and mitochondria) and cargo proteins (synapsin-1). Transport of cargo proteins, organelles, and different proteins from soma to axon terminals, along the length of axons, play an essential role in neuronal function ^37^. Therefore, our results suggests that excess accumulation of these cellular components in soma stresses cells, impairing neuronal function and eventually leading to increased ROS and cell death.

### TBCK knockdown alters mitochondrial quality in both proliferating and differentiated ReNcells

Anterograde and retrograde transport requires ATP, thus mitochondrial oxidative phosphorylation and mitochondrial-health play a critical role in neuronal homeostasis ^38^. Previous reports from TBCK patient fibroblast have compromised OXPHOS and mitochondrial biogenesis ^17^. To investigate if there is increased mitochondrial stress in TBCK KD ReNcells, and to determine readout mitochondrial function as an important readout for neurodegeneration, we stained NPC and differentiated cells with mitotraker dye. Recent evidence that TBCK is part of FERRY complex which carries number of mRNA that require for mitochondrial biogenesis ^20^, further suggests role mitochondrial dysfunction in TBCK KD ReNcells. When stained proliferating ReNcells with mitochondrial membrane-potential sensitive dye MitoTracker™ Orange CMTMRos, showed increased fragmentation and less dye accumulation in TBCK KD cells **(Figure. 7A)**. These results indicate that TBCK KD NPC have dysfunctional mitochondria. Further, we stained differentiated cells with the mitochondria marker ATP5B and the neuronal marker MAP2, and observed fewer mitochondria were present in the soma of TBCK KD neurons **(Figure. 7B)**. When we looked into the neuronal extensions, TBCK KD cells had overall less dense mitochondria distribution **(Figure. 7C)**. Additionally, we stained differentiated cells with MitoTracker™ Orange CMTMRos and found less dye accumulated in the soma **(Figure. 7D & E)**. Since we observed increased mitochondrial fragmentation in NPCs, we treated cells with mitochondrial fission inhibitors Mdivi-1 and observed a significant increase in the dye accumulation and improved mitochondrial network **(Figure. 7F & G)**, suggesting a partial rescue of this phenotype. These results indicate TBCK KD in ReNcells induces mitochondrial stress in both proliferating and differentiated cells. Furthermore, these results reveal a potential cause for cellular stress at NPC that leads to multiple cellular defects observed in differentiated TBCK KD cells.

**Figure 7.**
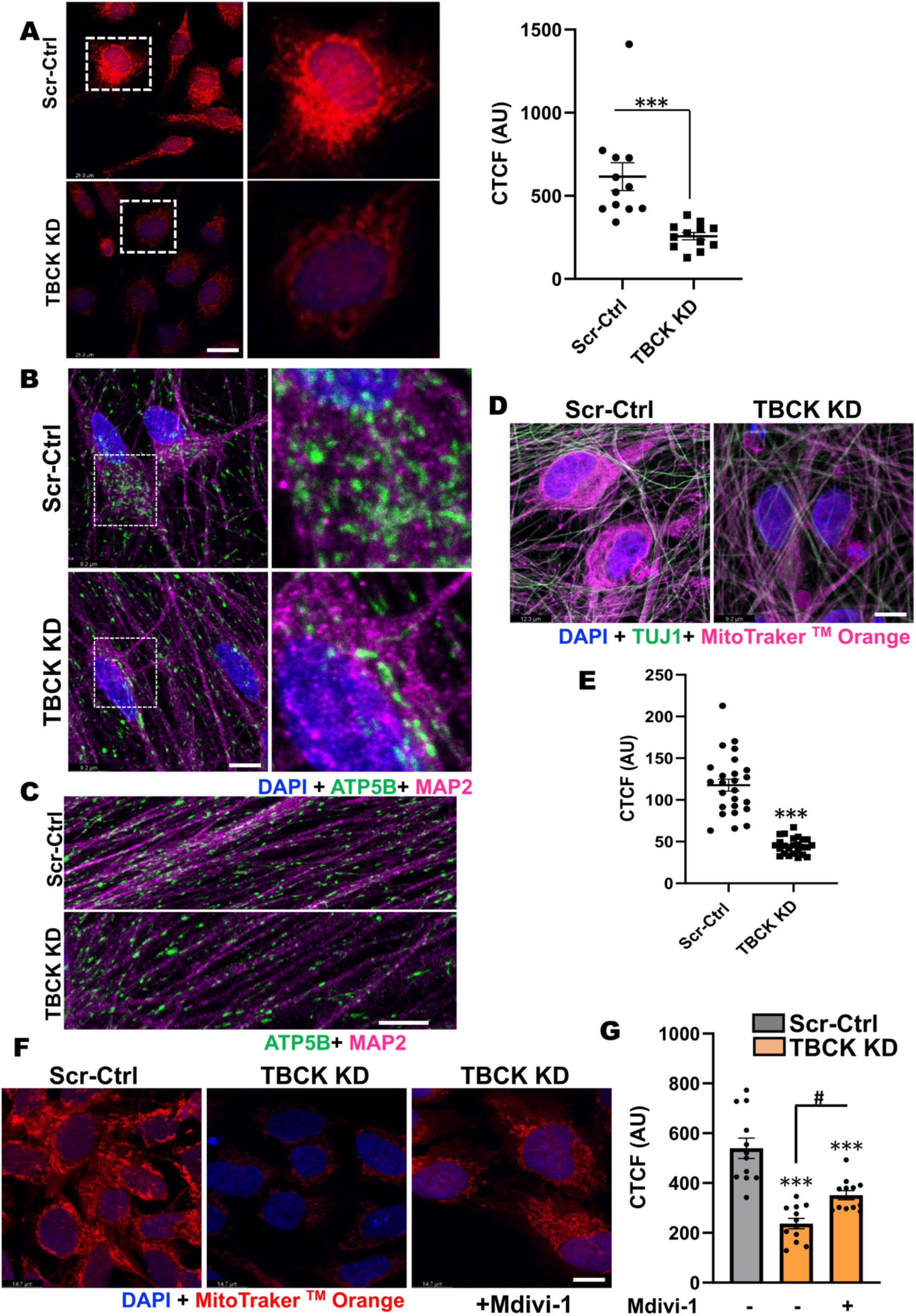
TBCK KD in ReNcells affected mitochondrial quality. **A:** Scr-Ctrl and TBCK KD proliferating ReNcells were treated with MitoTracker™ Orange CMTMRos and DAPI. Panels on the right show equally zoomed in views of panels on the left outlined in dashed-white boxes. Scale bar is 26.3µm. The fluorescence intensity of each cell is presented as CTCF. Error bars represent SEM. *** vs. Scr-Ctrl. **B & C: (B)** Immunofluorescence staining of ATP5B, MAP2, and DAPI for differentiated Scr-Ctrl and TBCK KD ReNcells. Panels on the right show equally zoomed in views of panels on the left outlined in dashed-white boxes. Images were taken at 100X magnification, scale bar is 9.2µm. **(C)** Enlarged images focused at neuronal extensions from Scr-Ctrl and TBCK KD cells were presented. **D:** Scr-Ctrl and TBCK KD differentiated ReNcells were treated with MitoTracker™ Orange CMTMRos and counter stained for TUJ1 and DAPI. Scale bar is 9.2µm. **E:** Scr-Ctrl and TBCK KD differentiated cells treated MitoTracker™ Orange CMTMRos and DAPI. The fluorescence intensity of each cell is presented as CTCF. Error bars represent SEM. ***vs. Control. **F:** TBCK KD proliferating ReNcells treated with Mdivi-1 shows rescue of MitoTracker^TM^ Orange phenotype loss observed in KD cells. Scale bar is 14.7µm. **G:** Quantification of Panel F showing significant decrease of MitoTracker™ Orange in TBCK KD cells and significant increase in TBCK KD cells after treatment with Mdivi-1. The fluorescence intensity of each cell is presented as CTCF. Error bars represent SEM. ***vs. Control, #vs. Mdivi-1.

## Discussion

Individuals diagnosed with TBCK syndrome show severe neurodegeneration, as evidenced by the progressive thinning of the corpus callosum and loss of brain volume observed in MRI scans ^2, 5, 16^. Many of these individuals exhibit progressive intellectual disability and seizures. However, prior to this publication cell-culture-based studies exploring TBCK function were solely performed in lymphoblasts or dermal fibroblasts. One recent study used iPSC derived NPC to understand the role of TBCK in vesicle secretion, and while this important study lacked isogenic controls and were not differentiated to neurons ^21^. To address whether loss of TBCK affects neurons, we generated a knockdown model in the well-characterized human NPC cells ReNcell VM, to provide insights into the role of TBCKs function in neurons. Our TBCK knockdown model revealed delays in neuronal and astrocyte maturation, likely due to reduced proliferation and increased cell death during or after differentiation. Our results are similar to Moreira et al., 2021 in that the NPC proliferation was affected. Further, we show that mitochondrial defects and altered gene expression in the proliferating NPC effected ReNcells differentiation into neurons and astrocytes. Interestingly, we observed decreased MAP2 (neuronal maturation marker) and increased GFAP (astrocyte marker). This could be due to either neurons dying or astrocytes over-proliferating. Based on our cumulative results, we believe the former due to neuronal dependence on mitochondrial oxidative phosphorylation, whereas astrocytes typically utilize glycolysis for their energy demands. Moreover, elevated autophagosome accumulation observed in TBCK-deficient neurons and astrocytes aligns with previous findings from TBCK patient fibroblasts ^16^. The observed decrease in the total amount of differentiated neurons coupled with increased apoptosis may indicate that individuals lacking functional copies of TBCK have a decreased capacity for neurodifferentiation and neuronal maturation, thus leading to neurodevelopmental delay and impaired neural function, and eventually neurodegeneration.

TBCK patients generally show phenotypes indicative of dysregulated function of the basal ganglia neural circuitry, including speech delay, seizures, poor psychomotor and musculoskeletal development ^2^. Dopaminergic (DA) neurons regulate voluntary movement, motivation, and habit learning as part of the basal ganglia circuitry in the midbrain ^32^. Their loss is associated with PD and AD ^33, 34^. When we differentiated ReNcells into DA neurons, the presence of fewer TH-positive staining and more neuronal extensions were broken, indicative of defects in DA neuronal function. These results are significant in that they suggest TBCK loss at NPC affects not only general cortical neuronal differentiation but specialized neurons, such as DA neurons, as well.

With respect to the molecular function of TBCK, our data provide new insights into the the role of mTORC1 in TBCK patients. Our current study varies from previous work showing universally decreased mTORC1 activity or no differences in activity ^21^, indicating that variation in signaling may exist between TBCK mutations, cell type, and growth factor presence. TBCK knockdown in ReNcells moderately increased mTORC1 activity both in NPCs and completly differentiated neurons. RNAseq results showed a number of mTORC1 signalling genes significantly upregulated in TBCK KD neurons and astrocytes. In eukaryotes, mTORC1 regulates cell growth and metabolism in response to nutrients, growth factors and stress. mTORC1 dysregulation is reported in AD/DS (Alzheimer’s, Down’s Syndrome), PD (Parkinson’s), HD (Huntington’s) and ALS (Amyotrophic Lateral Sclerosis) both in the brain and peripheral cells ^39^. Researchers observed both hyperactivation and hypoactivation in these neurodegenerative diseases, indicating that any dysregulation in mTORC1 affects brain function and viability. Although we observed moderately higher levels of phosphorylation of ribosomal protein S6 in TBCK KD ReNcells, future studies are required to fully understand how it may impact neuronal differentiation.

With our RNAseq screen, we used a high-sensitivity unbiased approach to look at comprehensive gene expression and uncovered several potential disease mechanisms by which TBCK regulates autophagy, progenitor proliferation, and neuronal differentiation. As expected based on our novel data, neuronal pathways were downregulated in ReNcells with targeted TBCK knockdown, including nervous system development, synaptic transmission, axon guidance, neurotransmitter receptor activity, neuron projections, postsynaptic density, glutamatergic and cholinergic synapses, neurexins, and neuroligins. These aligned well with the clinical phenotypes of seizures, brain atrophy, hypotonia, and respiratory deficiencies, supporting the hypothesis that the systemic deficits are secondary to neurologic dysfunction ^21, 40^. Interestingly, similar neurodegenerative pathways associated with Huntington’s disease were also up-regulated in TBCK KD cells, further emphasizing these cells as a good model for neurodegeneration in individuals with TBCK, and may also point to similar pathway dysregulation to those occurring in Huntington’s disease ^41^. On the other hand, epithelial-mesenchymal transition (EMT), a key process in development and wound healing, was also found to be up-regulated, although it is often associated with disease processes such as metastasis. In the context of TBCK, the significance of EMT upregulation is less clear, as none of the individuals with TBCK have reported early detection of cancer. However, these results could be pointing to a broader disruption in cell identity and function. For example, it is possible that TBCK is needed for the proliferation and differentiation of NPC during early stages of neurogenesis and thus contribute to neuronal apoptosis and axonal instability in TBCK-deficient or -mutated circumstances.

Of particular interest is the observed mitochondrial dysfunction in our model. As ATP is the primary energy sources for neurons, mitochondria play a critical role in supporting the high-energy demands associated with cargo transport, lipid membrane maintenance, and synaptic neurotransmission. Defects in mitochondrial function have previously been well documented to lead to neurodegenerative diseases including Parkinson’s Disease (PD), Alehimer’s Disease (AD), and Amyotrophic Lateral Sclerosis ^42, 43, 44^. Several mitochondria-related pathways have previously been associated with cell stress, and were found to be up-regulated in our TBCK KD cells. These included NO synthase regulation, ATP synthesis coupled with electron transport, mitochondrial elongation and termination, glutathione metabolism, and oxidative phosphorylation. As higher energetic demands are placed on cells, these pathways commonly are called upon to help achieve cell homeostasis. Of particular note was the up-regulation of ferroptosis as well as several biosynthetic processes related to lipid membrane metabolism. Ferroptosis is a unique form of regulated cell death characterized by iron-dependent lipid peroxidation ^45^ and showed significant upregulation in TBCK knockdown cells. Abnormal regulation of ferroptosis has also been implicated in a variety of diseases that have neurodegeneration, including Friedrich’s Ataxia, PD, AD, and Huntington’s Disease (HD) ^46^. Specifically, glutathione homeostasis is one of the main metabolic pathways that regulate ferroptosis induction. If unchecked, ferroptosis can lead to neuronal death, thus possibly contributing to the pathogenesis of neurodegeneration observed in TBCK syndrome. Increased mitochondrial fragmentation and partial rescue with the mitochondiral fission inhibior mdivi1 suggests that TBCK is necessary for mitochondrial function. Mitochondrial defects in NPCs, and improving mitochondrial dynamics with fission inhibitors implicate that this pathway is a potential treatment modality to improve neuronal maturation and survival of these cells.

The accumulation of autophagosomes and transport proteins seen in our TBCK-deficient model may also be indicative of heightened cellular stress, leading to premature cell death and defects in neuronal differentiation observed in our model. This is particularly relevant given that various neurodegenerative diseases like PD, AD, HD, and ALS have specific mutated proteins that accumulate causing cytotoxicity that contribute to disease pathogenesis ^47, 48, 49^. Previous studies of TBCK patient-derived fibroblasts have reported a higher accumulation of LC3 positive vesicles that is likely due to incomplete degradation of autophagosomes inside the lysosomes ^16^. Our study with HeLa cells expressing LC3 protein conjugated with GFP and RFP proved that lysosomal degradation of autophagosomes is the primary reason for increased autophagy. While more evidence is needed, these findings support TBCK’s role as an important mediator for mitochondrial homeostasis and autophagy.

Taken together, our findings indicate that NPC proliferation and neuronal maturation are central to the pathogenesis in children with TBCK syndrome. Specifically, recent studies have identified TBCK as a part of a multiprotein complex called the FERRY ^20, 50^, which transport mRNA to different parts of cell using RAB5 and early endosomal pathway. This cellular process is essential in humans, who harbor long axonal tracts where local protein translation plays an essential role in orgenelle biosynthesis. Therefore, the proposed TBCK FERRY complex likely plays an essential role in neuronal health by supporting local protein translation in these axonal tracts. While our studies support a function of TBCK in protein translation and transport, further analyses are needed to determine whether or not these are direct causes of neurodegeneration. Furthermore, our model reveals a severe reduction of neurite projection, likely impacting axonal tract formation. We also observed dysregulation in the EEA1 distribution of the differentiated cells, and when considered with Moreira et al.’s findings of aberrant Clathrin, initiation of endocytosis may be considered for the role of the TBCK protein ^21^. Therefore, TBCK’s role in axonal transport may be secondarily important compared to other defects illustrated in our study. TBCK syndrome is a severe and multi-organ dysfunction disease, and further studies are required to understand the role of TBCK in different organ systems. Although we showed partial rescue with the fission inhibitor, systemic studies using appropriate cell models are needed to fully undestand how this treatment can rescue transport dysfunction in other organ systems. In summary, our ReNcell model of TBCK knockdown provides novel potential pathogenic cellular mechanisms of disease, which open new avenues for therapeutic intervention both in early development and postnatally.

## Supporting information

differentially expressed genes and pathways

## Acknowledgements

Thank you to the Ortiz-Gonzalez laboratory at the Children’s Hospital of Philadelphia for sharing their TBCK patient-derived dermal fibroblast lines.

## References

1. Zapata-Aldana E, Kim DD, Remtulla S, Prasad C, Nguyen CT, Campbell C. Further delineation of TBCK - Infantile hypotonia with psychomotor retardation and characteristic facies type 3. Eur J Med Genet 62, 273–277 (2019).

2. Bhoj EJ, et al. Mutations in TBCK, Encoding TBC1-Domain-Containing Kinase, Lead to a Recognizable Syndrome of Intellectual Disability and Hypotonia. Am J Hum Genet 98, 782–788 (2016).

3. Durham EL, et al. TBCK syndrome: a rare multi-organ neurodegenerative disease. Trends Mol Med, (2023).

4. Beck-Wodl S, et al. Homozygous TBC1 domain-containing kinase (TBCK) mutation causes a novel lysosomal storage disease - a new type of neuronal ceroid lipofuscinosis (CLN15)? Acta Neuropathol Commun 6, 145 (2018).

5. Chong JX, et al. Recessive Inactivating Mutations in TBCK, Encoding a Rab GTPase-Activating Protein, Cause Severe Infantile Syndromic Encephalopathy. Am J Hum Genet 98, 772–781 (2016).

6. Wu J, Lu G. Multiple functions of TBCK protein in neurodevelopment disorders and tumors. Oncol Lett 21, 17 (2021).

7. Pan X, Eathiraj S, Munson M, Lambright DG. TBC-domain GAPs for Rab GTPases accelerate GTP hydrolysis by a dual-finger mechanism. Nature 442, 303–306 (2006).

8. Stenmark H, Olkkonen VM. The Rab GTPase family. Genome Biol 2, REVIEWS3007 (2001).

9. Veleri S, Punnakkal P, Dunbar GL, Maiti P. Molecular Insights into the Roles of Rab Proteins in Intracellular Dynamics and Neurodegenerative Diseases. Neuromolecular Med 20, 18–36 (2018).

10. Kiral FR, Kohrs FE, Jin EJ, Hiesinger PR. Rab GTPases and Membrane Trafficking in Neurodegeneration. Curr Biol 28, R471–R486 (2018).

11. Liu Y, Yan X, Zhou T. TBCK influences cell proliferation, cell size and mTOR signaling pathway. PLoS One 8, e71349 (2013).

12. Saxton RA, Sabatini DM. mTOR Signaling in Growth, Metabolism, and Disease. Cell 169, 361–371 (2017).

13. LiCausi F, Hartman NW. Role of mTOR Complexes in Neurogenesis. Int J Mol Sci 19, (2018).

14. Lamming DW, Sabatini DM. A Central role for mTOR in lipid homeostasis. Cell Metab 18, 465–469 (2013).

15. Mao Z, Zhang W. Role of mTOR in Glucose and Lipid Metabolism. Int J Mol Sci 19, (2018).

16. Ortiz-Gonzalez XR, et al. Homozygous boricua TBCK mutation causes neurodegeneration and aberrant autophagy. Ann Neurol 83, 153–165 (2018).

17. Tintos-Hernandez JA, Santana A, Keller KN, Ortiz-Gonzalez XR. Lysosomal dysfunction impairs mitochondrial quality control and is associated with neurodegeneration in TBCK encephaloneuronopathy. Brain Commun 3, fcab215 (2021).

18. Rangaraju V, et al. Pleiotropic Mitochondria: The Influence of Mitochondria on Neuronal Development and Disease. J Neurosci 39, 8200–8208 (2019).

19. Kann O, Kovacs R. Mitochondria and neuronal activity. Am J Physiol Cell Physiol 292, C641–657 (2007).

20. Schuhmacher JS, et al. The Rab5 effector FERRY links early endosomes with mRNA localization. Mol Cell 83, 1839–1855 e1813 (2023).

21. Moreira DP, et al. Neuroprogenitor Cells From Patients With TBCK Encephalopathy Suggest Deregulation of Early Secretory Vesicle Transport. Front Cell Neurosci 15, 803302 (2021).

22. Donato R, et al. Differential development of neuronal physiological responsiveness in two human neural stem cell lines. BMC Neurosci 8, 36 (2007).

23. Ortinau S, et al. Effect of 3D-scaffold formation on differentiation and survival in human neural progenitor cells. Biomed Eng Online 9, 70 (2010).

24. Bridges CR, et al. USP9X deubiquitylating enzyme maintains RAPTOR protein levels, mTORC1 signalling and proliferation in neural progenitors. Sci Rep 7, 391 (2017).

25. Dobin A, et al. STAR: ultrafast universal RNA-seq aligner. Bioinformatics 29, 15–21 (2013).

26. Li B, Dewey CN. RSEM: accurate transcript quantification from RNA-Seq data with or without a reference genome. BMC Bioinformatics 12, 323 (2011).

27. Love MI, Huber W, Anders S. Moderated estimation of fold change and dispersion for RNA-seq data with DESeq2. Genome Biol 15, 550 (2014).

28. Tarazona S, Furió-Tarı P, Ferrer A, Conesa A. NOISeq: Differential Expression in RNA-seq. Version 216, (2013).

29. Conway JR, Lex A, Gehlenborg N. UpSetR: an R package for the visualization of intersecting sets and their properties. Bioinformatics 33, 2938–2940 (2017).

30. Gu Z, Eils R, Schlesner M. Complex heatmaps reveal patterns and correlations in multidimensional genomic data. Bioinformatics 32, 2847–2849 (2016).

31. 31. Loos B, du Toit A, Hofmeyr JH. Defining and measuring autophagosome flux-concept and reality. Autophagy 10, 2087–2096 (2014).

32. Luo SX, Huang EJ. Dopaminergic Neurons and Brain Reward Pathways: From Neurogenesis to Circuit Assembly. Am J Pathol 186, 478–488 (2016).

33. Surmeier DJ. Determinants of dopaminergic neuron loss in Parkinson’s disease. FEBS J 285, 3657–3668 (2018).

34. Nobili A, et al. Dopamine neuronal loss contributes to memory and reward dysfunction in a model of Alzheimer’s disease. Nat Commun 8, 14727 (2017).

35. Yang WS, Stockwell BR. Ferroptosis: Death by Lipid Peroxidation. Trends Cell Biol 26, 165–176 (2016).

36. Hilfiker S, Pieribone VA, Czernik AJ, Kao HT, Augustine GJ, Greengard P. Synapsins as regulators of neurotransmitter release. Philos Trans R Soc Lond B Biol Sci 354, 269–279 (1999).

37. Sleigh JN, Rossor AM, Fellows AD, Tosolini AP, Schiavo G. Axonal transport and neurological disease. Nat Rev Neurol 15, 691–703 (2019).

38. Yellen G. Fueling thought: Management of glycolysis and oxidative phosphorylation in neuronal metabolism. J Cell Biol 217, 2235–2246 (2018).

39. Querfurth H, Lee HK. Mammalian/mechanistic target of rapamycin (mTOR) complexes in neurodegeneration. Mol Neurodegener 16, 44 (2021).

40. Murdock DR, et al. Transcriptome-directed analysis for Mendelian disease diagnosis overcomes limitations of conventional genomic testing. J Clin Invest 131, (2021).

41. Li S, Li XJ. Multiple pathways contribute to the pathogenesis of Huntington disease. Mol Neurodegener 1, 19 (2006).

42. Trigo D, Avelar C, Fernandes M, Sa J, da Cruz ESO. Mitochondria, energy, and metabolism in neuronal health and disease. FEBS Lett 596, 1095–1110 (2022).

43. Lin MT, Beal MF. Mitochondrial dysfunction and oxidative stress in neurodegenerative diseases. Nature 443, 787–795 (2006).

44. Lin F, Luo SQ. Mitochondria in neurodegenerative diseases. CNS Neurosci Ther 25, 813–815 (2019).

45. Maher P, Currais A, Schubert D. Using the Oxytosis/Ferroptosis Pathway to Understand and Treat Age-Associated Neurodegenerative Diseases. Cell Chem Biol 27, 1456–1471 (2020).

46. Kim SW, Kim Y, Kim SE, An JY. Ferroptosis-Related Genes in Neurodevelopment and Central Nervous System. Biology (Basel) 10, (2021).

47. Fleming A, et al. The different autophagy degradation pathways and neurodegeneration. Neuron 110, 935–966 (2022).

48. Corti O, Blomgren K, Poletti A, Beart PM. Autophagy in neurodegeneration: New insights underpinning therapy for neurological diseases. J Neurochem 154, 354–371 (2020).

49. Frake RA, Ricketts T, Menzies FM, Rubinsztein DC. Autophagy and neurodegeneration. J Clin Invest 125, 65–74 (2015).

50. Quentin D, et al. Structural basis of mRNA binding by the human FERRY Rab5 effector complex. Mol Cell 83, 1856–1871 e1859 (2023).

