## Supplementary tables and figures for "A Novel Human TBCK- Neuronal Cell Model Results in Severe Neurodegeneration and Partial Rescue with Mitochondrial Fission Inhibition"

### Supplementary Figure. 1

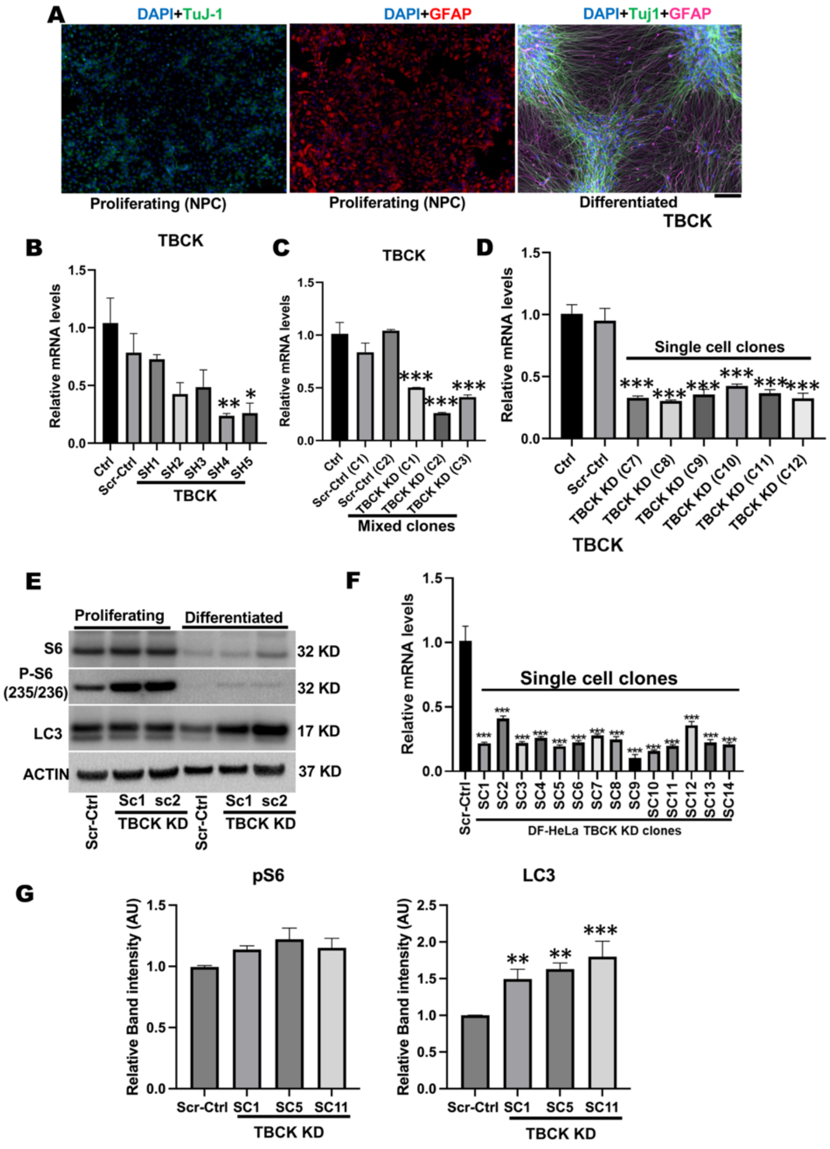


**(A)**Proliferating and differentiated ReNcells were stained with DAPI, TUJ1, and GFAP antibodies. Images were taken at 20X magnification. Images are representation of three independent experiments. **(B)** qRT-PCR analysis of TBCK mRNA levels in 293T cells transfected with 2µg of different pLKO.1 plasmid with different shRNA targeting TBCK. *vs Control. **(C)** qRT-PCR analysis of TBCK mRNA levels in ReNcells transduced with different dilutions of lentivirus. Lentivirus transduced clones #1-3 significantly reduced TBCK mRNA levels compared to Scr-ctrl. *vs Control. **(D)** qRT-PCR analysis of TBCK mRNA levels in lentivirus-transduced single-cell clones. Knockdown in clones #7-12 shows a significant reduction in TBCK mRNA levels compared to controls. *vs Control.**(E)**Western blots of proliferating and differentiated ReNcells for S6, p-S6, LC3, and actin. WB represents three independent experiments. **(F)** qRT-PCR analysis of TBCK mRNA levels in lentivirus transduced single cell clones of DF-HeLA cells. *vs Scr-Ctrl. **(G)** Western blot quantifications of Scr-Ctrl and TBCK KD DF-HeLa clones for pS6 and LC3-II. Band intensities were calculated using Image J normalized to Actin. The relative band intensity compared to the Scr-Ctrl was presented. Error bars represent SEM. vs. Control. *vs Scr-Ctrl (*p-value < 0.05, **p-value < 0.01, ***p-value < 0.001).

**Supplementary Figure. 2**

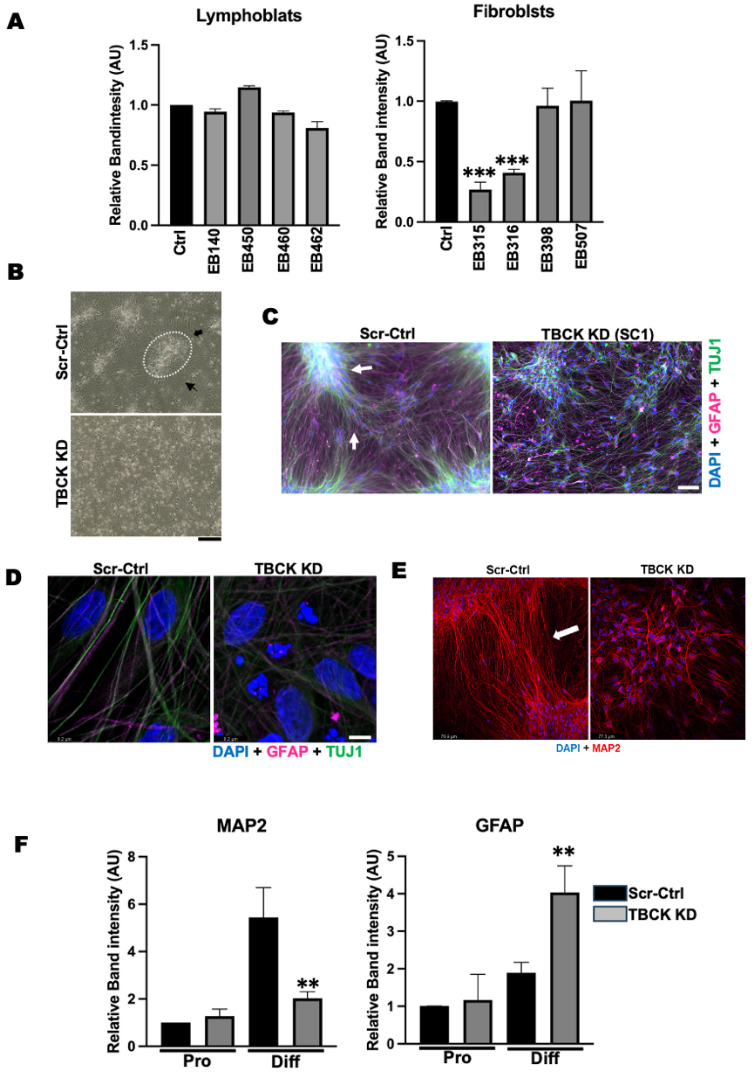


**(A)** Western blot quantifications of controls and patent-derived lymphoblast and fibroblasts for pS6 levels. Band intensities were calculated using Image J normalized to Actin. The relative band intensity compared to the Control was presented. *vs. Control. (**B)** Scr-Ctrl and TBCK KD ReNcells were differentiated into neurons and astrocytes and light microscope images were taken using EVOS XL microscope at 10X magnification. Images represent three independent experiments. **(C)** Immunofluorescence staining of TUJ1, GFAP, and DAPI for differentiated Scr-Ctrl and TBCK KD-sc1 ReNcells. Images were taken at 20X magnification. Images represent three independent experiments. **(D)** Immunofluorescence staining of TUJ1, GFAP, and DAPI for differentiated Scr-Ctrl and TBCK KD ReNcells. Images were taken at 63X magnification and zoomed into 4X. Images represent three independent experiments. **(E)** Immunofluorescence staining of MAP2 and DAPI for differentiated Scr-Ctrl and TBCK KD ReNcells. Images were taken at 20X magnification. Images represent three independent experiments. **(F)** Western blot quantifications of proliferating and differentiated ReNcells for MAP2 and GFAP levels. Band intensities were calculated using Image J normalized to Actin. The relative band intensity compared to the proliferating Scr-Ctrl was presented. *vs. Differentiated Scr-Ctrl. (*p-value < 0.05, **p-value < 0.01, ***p-value < 0.001).

### Supplementary Figure. 3

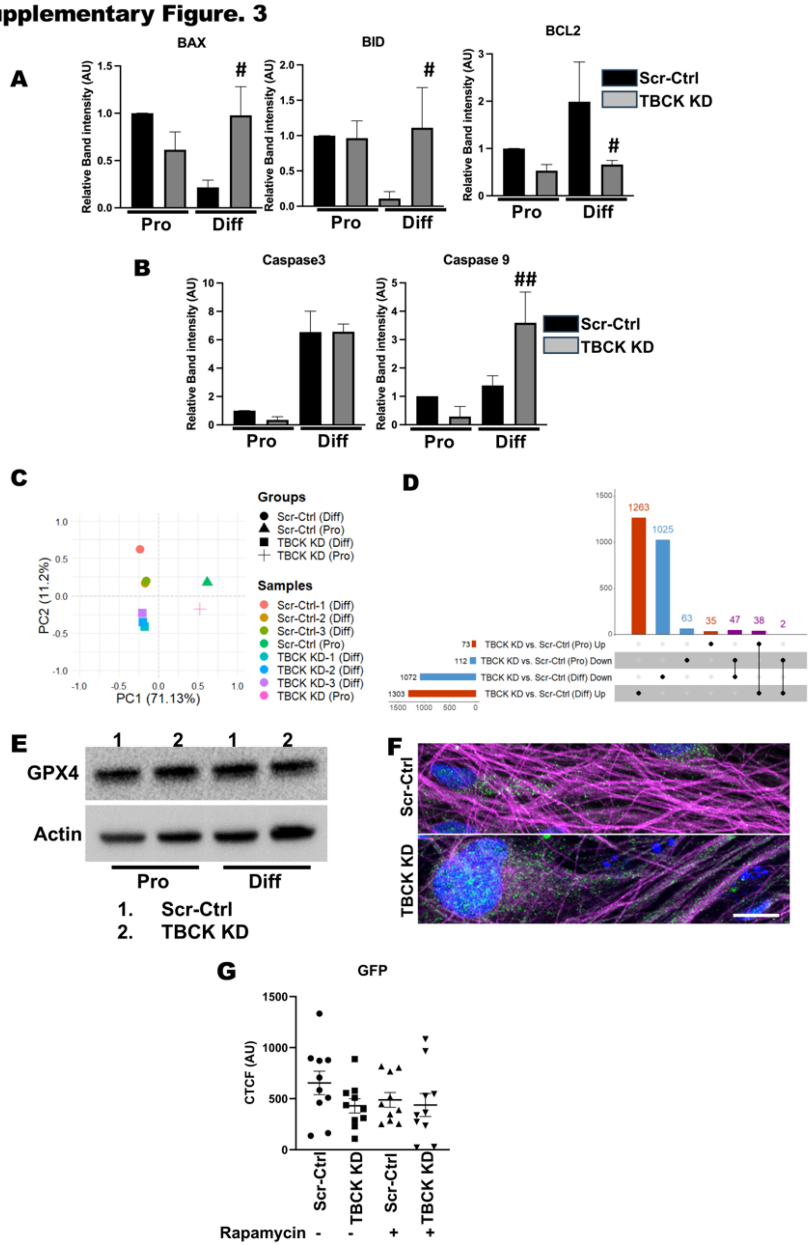


(A)Western blot quantifications of proliferating and differentiated ReNcells for BAX, BID, and BCL2 levels. (B)Western blot quantifications of proliferating and differentiated ReNcells for Caspase 3 and 9 levels. Band intensities were calculated using Image J normalized to Actin. The relative band intensity compared to the proliferating Scr-Ctrl was presented. #vs. Differentiated Scr-Ctrl. (#p-value < 0.05, ##p-value < 0.01). (C)Principal component analysis (PCA) proliferating and differentiated cells of both Control and KD groups. Shape represents the groups, and color represents the samples. Distinct clusters of samples indicate the effect of cell stage and KD. (D)The upSet plot represents the distribution and intersection of differentially expressed genes between Proliferating cells: KD vs. Control and Differentiated cells: KD vs. Control. Rows represent the number of peaks in each comparison (Proliferating cells: KD vs. Control and Differentiated cells: KD vs. Control), and columns represent the number of genes per interaction. The red and blue horizontal bars represent the increased and decreased gene expression, respectively. The black filled dots indicate the association between rows. The red and blue vertical bars represent genes unique to a given comparison, while the vertical purple bars represent the number of genes overlap between groups.(E)Western blots of proliferating and differentiated ReNcells for GPX4 and actin. WB represents three independent experiments.(F)Immunofluorescence staining of LC3, TUJ1, and DAPI for differentiated Scr-Ctrl and TBCK KD ReNcells. Images were taken at 63X magnification. (G)Scr-Ctrl and TBCK KD DF-HeLa cells treated with rapamycin and images were taken at 63X magnification. The GFP intensities of the cells were calculated using image J and expressed as corrected total cell fluorescence (CTCF).

### Supplementary Figure. 4

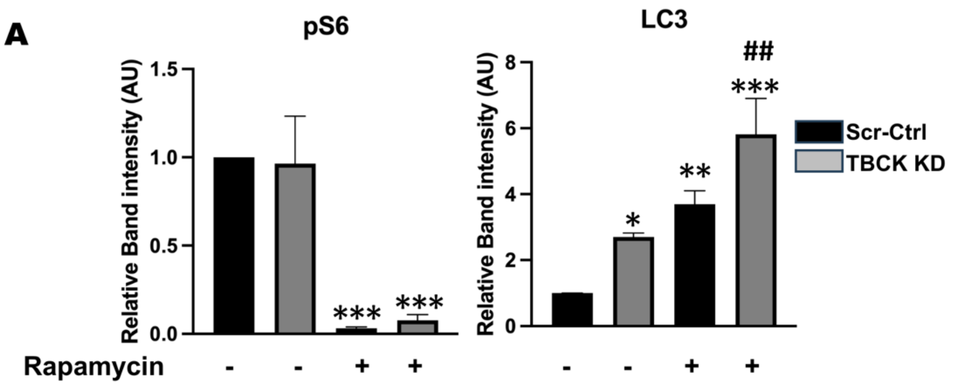


(**A**) Western blot quantifications of Scr-Ctrl and TBCK KD DF-HeLa cells treated with rapamycin for pS6 and LC3-II. Band intensities were calculated using Image J normalized to Actin. The relative band intensity compared to the untreated Scr-Ctrl was presented. *vs. Untreated Scr-Ctrl and #vs Rapamycin treated Scr-Ctrl. (*p-value < 0.05, **p-value < 0.01, ***p-value < 0.001).

### Supplementary Tables:

**Table.1** Different patient-derived cells with differing TBCK mutations presented as a table.

| Patient ID | Cell line | Clinical Information |
|---|---|---|
| EB140 | Lymphoblast | Compound heterozygous mutation (loss of function) |
| EB316 | Fibroblast | Homozygous loss function mutation |
| EB462 | Lymphoblast | Compound heterozygous mutation (loss of function) |
| EB507 | Fibroblast | Compound heterozygous mutation (loss of function) |
| EB315 | Fibroblast | Homozygous loss function mutation |
| EB398 | Fibroblast | Homozygous loss function mutation |
| EB450 | Lymphoblast | Homozygous loss function mutation |
| EB460 | Lymphoblast | Homozygous loss function mutation |

**Table. 2** Primers and their sequences used in the present study.

| | |
|---|---|
| **TBCK-F.P** | **TGAAGAGCATGGTTGTTTGGAC** |
| **TBCK-R.P** | **GAAGGATGGAAGGTAAGGCAC** |
| **GAPDH-F.P** | **TGTGGGCATCAATGGATTTGG** |
| **GAPDH-R.P** | **ACACCATGTATTCCGGGTCAAT** |
| **MAP2-F.P** | **AGGCTGTAGCAGTCCTGAAAGG** |
| **MAP2-R.P** | **CTTCCTCCACTGTGACAGTCTG** |
| **GFAP-F.P** | **CTGGAGAGGAAGATTGAGTCGC** |
| **GFAP-R.P** | **ACGTCAAGCTCCACATGGACCT** |
| **TBCK-SH5** | **GACAAATTGAAGTGGATATTC** |
| **TBCK-SH4** | **AGAAGTTCGGCACCTTATTTC** |
| **TBCK-SH3** | **GCAAGAAACTTGACTCTACAT** |
| **TBCK-SH2** | **GCTAAAGGCTTATCCATATAA** |
| **TBCK-SH1** | **CAGTGAAATCTGATCATATAT** |

**Table. 3** Antibodies and their dilutions

| Antibody (Dilution used in the study) | Company and Catalogue number |
|---|---|
| S6 (1:1000) | 2317S (cell signaling) |
| p-S6 (1:1000) | 2211S (cell signaling) |
| PAX6 (1:1000) | PB9768 (Boster Bio) |
| MAP2 (1:1000) | 4542S (cell signaling) |
| GFAP (1:2000) | NB120-16997 (Novus) |
| TUJ1 (1:200) | ML1195Z (R&D systems) |
| SYN1 (1:100) | NB300-104 (Novus) |
| LAMP1 (1:100) | SC-20011 (Santa Cruz) |
| ACTIN (1:2000) | SC-69879 (Santa Cruz) |
| BrdU (1:100) | MA3-071 (Invitrogen) |
| GPX4 (1:1000) | 52455S (cell signaling) |
| Anti-Mouse Alexa 488 (1:500) | A11001 (Invitrogen) |
| Anti-Mouse Alexa 594 (1:500) | A32742 (Invitrogen) |
| Anti-Rabbit Alexa 488 (1:500) | SAB4600044 (Sigma) |
| Anti-Rabbit Alexa 594 (1:500) | A11037 (Invitrogen) |
